# Multisensory coding of audiovisual movies in the human hippocampus

**DOI:** 10.64898/2026.03.01.708855

**Authors:** Omri Raccah, Aryan Agarwal, Yannan Zhu, Nicholas B. Turk-Browne

## Abstract

The hippocampus receives convergent input from multiple sensory systems, yet in humans it has been studied almost exclusively through vision. Here we examine how the hippocampus contributes to sensory processing beyond the visual modality and to the multisensory integration of visual information with these other modalities. Participants were exposed to multiple repetitions of short naturalistic movie clips, each presented in four formats: auditory-only, visual-only, congruent audiovisual, and incongruent audiovisual (audio and video from different movies). Using high-resolution fMRI, we measured univariate activation and multivariate representations across the longitudinal axis and subfields of the human hippocampus. Whereas univariate analyses detected only visual responses across hippocampal subregions, with no activation for auditory stimuli and no benefit of congruent stimuli, multivariate analyses revealed robust representations of both auditory and visual scenes. The posterior hippocampus showed enhanced pattern similarity for congruent stimuli relative to unisensory stimuli, especially in the left hemisphere, demonstrating multisensory facilitation. The anterior hippocampus showed crossmodal decoding between auditory and visual versions of the same clip, driven by the right hemisphere, suggesting a more abstract representation. Finally, whole-brain searchlight analyses revealed parallel effects in cortical regions known to support multisensory integration. These findings advance understanding of auditory and multisensory coding in the human hippocampus, revealing specialization and lateralization along its longitudinal axis.

## Introduction

The hippocampus and surrounding medial temporal lobe (MTL) cortex receive anatomical inputs from virtually all sensory systems [1]. In non-human animals, hippocampal neurons encode auditory, tactile, olfactory, and visual information, and represent crossmodal associations [2, 3, 4]. For example, macaque hippocampal neurons exhibit selectivity for congruent face and voice pairings of individual animals [4]. Yet, in humans, the hippocampus has been studied almost exclusively in the visual modality [5]. This leaves a fundamental gap: How does the human hippocampus represent sensory modalities beyond vision, and how does it integrate sense information across modalities? Understanding this integration is fundamental to explaining how the brain constructs coherent experiences and memories from fragmented inputs.

Multisensory learning enhances memory across species, from Drosophila to humans [6, 7, 8, 9]. However, whether hippocampal mechanisms support this mnemonic benefit remains unknown. Determining the contribution of the hippocampus to multisensory processing requires considering its broader representational capacity. Beyond its traditional role in memory, an emerging literature suggests that the human hippocampus contributes to central aspects of visual perception [10, 11], including eye movements [12, 13], imagery [14, 15], scene discrimination [16, 17], and visual attention [18, 19]. Given that the hippocampus represents complex conjunctions of visual features, it may additionally be critical for binding information across sensory modalities.

We tested this possibility for auditory and visual modalities. In line with recent approaches [11], we focused on characterizing representational coding schemes in the hippocampus, as opposed to testing a specific role in perception or memory behavior. Using high-resolution fMRI in 30 participants, we examined unisensory and multisensory processing in the whole hippocampus, along its longitudinal axis, across hemispheres, and within its subfields. Participants were exposed to multiple repetitions of short naturalistic movie clips presented in four formats: auditory-only, visual-only, congruent audiovisual, and incongruent audiovisual (audio and video from different clips). We first conducted a general linear model (GLM) analysis to estimate univariate activation in each condition across hippocampal ROIs. Next, we performed representational similarity analysis (RSA) to investigate multivariate representations of unimodal and multimodal stimuli in these regions. We repeated these analyses across the whole brain to juxtapose effects in hippocampal ROIs with known cortical sites supporting multisensory integration.

This approach allows us to address three questions in this study: First, how is auditory information distributed in the hippocampus? Whereas auditory processing in the hippocampus has been widely studied in other species, evidence in humans remains limited (for review, see [5]). Only a handful of human fMRI studies have directly examined auditory processing in the human hippocampus [20, 21], and to our knowledge, none have mapped these representations across its anatomical subdivisions. Second, how does congruent multisensory stimulation facilitate processing in the hippocampus? Decades of research has demonstrated that congruent audiovisual stimuli enhance neural responses relative to unisensory stimuli, effects most prominent in the superior temporal sulcus (STS) and posterior superior temporal gyrus (posterior STG; e.g., [22, 23, 24, 25]). Whether the hippocampus exhibits similar multisensory facilitation effects remains unknown. Third, what is the relationship between auditory and visual representations in the hippocampus? The anterior subdivision of the hippocampus has been shown to maintain abstract (as opposed to perceptually detailed) representations in the visual modality [26, 27]. This raises the intriguing possibility that the anterior hippocampus may support generalization across sensory modalities that enables crossmodal transfer.

## Results

### Behavior

Participants first viewed 10 short movie clips in their original audiovisual format in the scanner prior to data acquisition (6 s each; Figure 1A). This pre-exposure was intended to limit memory demands in the main experiment by reducing novelty and learning, consistent with prior work that isolated representational content in the hippocampus [11].

**Figure 1:**
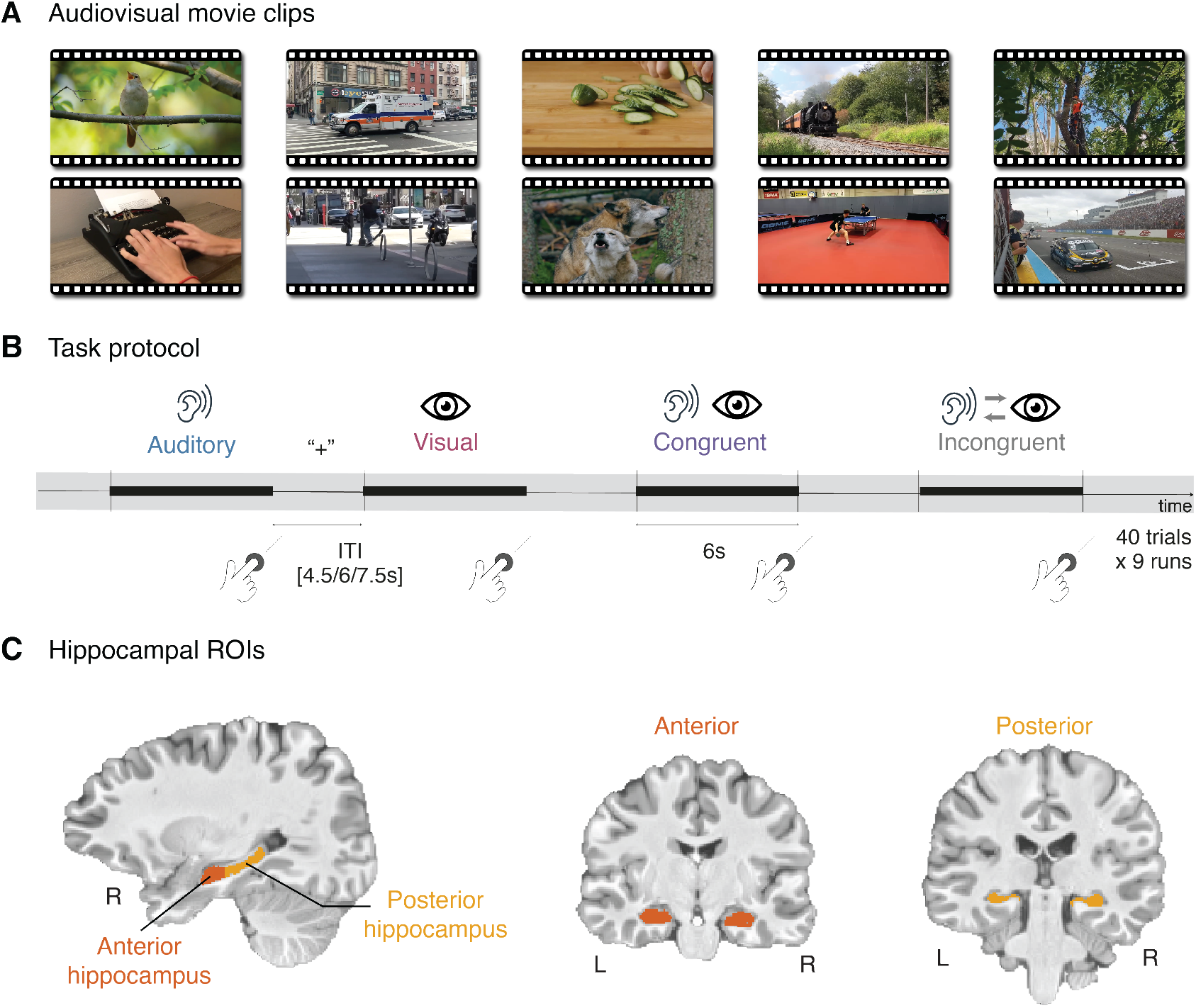
Experimental design and hippocampal segmentation. **(A)** The stimuli consisted of 10 naturalistic audiovisual movie clips (6 s each). **(B)** Unisensory stimuli were created by isolating the audio or visual track of each movie, resulting in 10 auditory-only and 10 visual-only stimuli. Incongruent audiovisual stimuli were generated by pairing audio and video tracks from different movies (90 unique pairings). In each run (9 runs total), all 10 stimuli from the auditory-only, visual-only, and congruent audiovisual conditions were presented, along with 10 incongruent audiovisual stimuli (40 total trials per run). Stimulus order was randomized within each run, with jittered inter-trial intervals between trials. As a cover task to ensure attention to the stimuli, participants were instructed to press a button within 1 s of each movie’s offset. **(C)** The hippocampal long axis was manually segmented into anterior and posterior subdivisions based on the last coronal slice in which the uncus was visible [28]. Example coronal slices through the anterior and posterior hippocampus.

During the main experiment, participants were exposed to multiple repetitions of these movie clips across four presentation formats: auditory-only, visual-only, congruent audiovisual, and incongruent audiovisual (mismatched audio and video from different clips; Figure 1B). All 4 formats of each of the 10 movies were presented once per run (40 stimuli per run) across nine runs, enabling leave-one-run-out cross-validation analyses. For the incongruent audiovisual condition, each video was paired once with audio from each of the other movies; these re-pairings varied across runs to ensure that the visual and auditory content of each movie appeared as part of an incongruent trial once per run. A central fixation cross remained on screen throughout each 6-s trial across all conditions, superimposed over videos in the visual-only and audiovisual conditions.

To ensure attention during the movies, participants were instructed to press a button within 1 s of each movie’s offset (Figure 1B). A response was scored as correct if it fell within this 1-s post-offset window, whereas presses during the movie or after the window were scored as incorrect. Performance in this offset detection task was reliable and near ceiling in every condition (auditory: mean accuracy across participants, *M* = 0.94, standard error of the mean, *SEM* = 0.006; visual: *M* = 0.98, *SEM* = 0.006; congruent: *M* = 0.98, *SEM* = 0.003; incongruent: *M* = 0.99, *SEM* = 0.003). Response times (RTs) on correct trials were fast and consistent across conditions (auditory: *M* = 379 ms, *SEM* = 13; visual: *M* = 370 ms, *SEM* = 13; congruent: *M* = 337 ms, *SEM* = 11; incongruent: *M* = 334 ms, *SEM* = 10), confirming participants’ engagement throughout the experiment.

### Univariate activity

To examine univariate activity, we constructed a GLM with regressors for each condition. The resulting beta estimates were extracted from each region of interest (ROI) and averaged across voxels per participant. We focused ROI analyses on the whole hippocampus, anterior and posterior hippocampus, the left and right hippocampus, and hippocampal subfields CA1, CA2/3, DG, and subiculum. We examined subfields given evidence for differential audiovisual selectivity across subfields in the primate hippocampus [4]. We also examined MTL cortex, comprising the parahippocampal cortex (PHC), perirhinal cortex (PRC), and entorhinal cortex (EC). We corrected for multiple comparisons separately across anterior/posterior subdivisions, hemispheres, subfields, and MTL cortex.

The whole hippocampus (Figure 2A) showed significant evoked responses to unisensory visual stimuli (*P <* 0.001) and both multisensory audiovisual conditions (congruent: *P <* 0.001; incongruent: *P <* 0.001), but not to unisensory auditory stimuli (*P* = 0.55). This pattern was consistent in the anterior hippocampus (visual: *P <* 0.001; auditory: *P* = 0.89; congruent: *P <* 0.001; incongruent: *P* = 0.001) and posterior hippocampus (visual: *P <* 0.001; auditory: *P* = 0.42; congruent: *P <* 0.001; incongruent: *P <* 0.001), as well as in the left hippocampus (visual: *P <* 0.001; auditory: *P* = 0.75; congruent: *P <* 0.001; incongruent: *P <* 0.001) and the right hippocampus (visual: *P <* 0.001; auditory: *P* = 0.75; congruent: *P <* 0.001; incongruent: *P <* 0.001; Figure 2B). All hippocampal subfields showed this same profile, with significant responses to the visual and both audiovisual conditions but no detectable response to auditory-only stimuli (Figure S1B). We found no evidence of multisensory facilitation in univariate activity: the congruent audiovisual condition did not reliably exceed the visual condition in any hippocampal ROI (all *P >* 0.05). There was a marginal trend of suppression for the incongruent condition relative to the congruent audiovisual condition in the whole hippocampus, its left and right hemispheres, and the anterior hippocampus (all *P <* 0.1). For full descriptive statistics and results from sensory and MTL cortical ROIs, see Table S1 and Figure S2.

**Figure 2:**
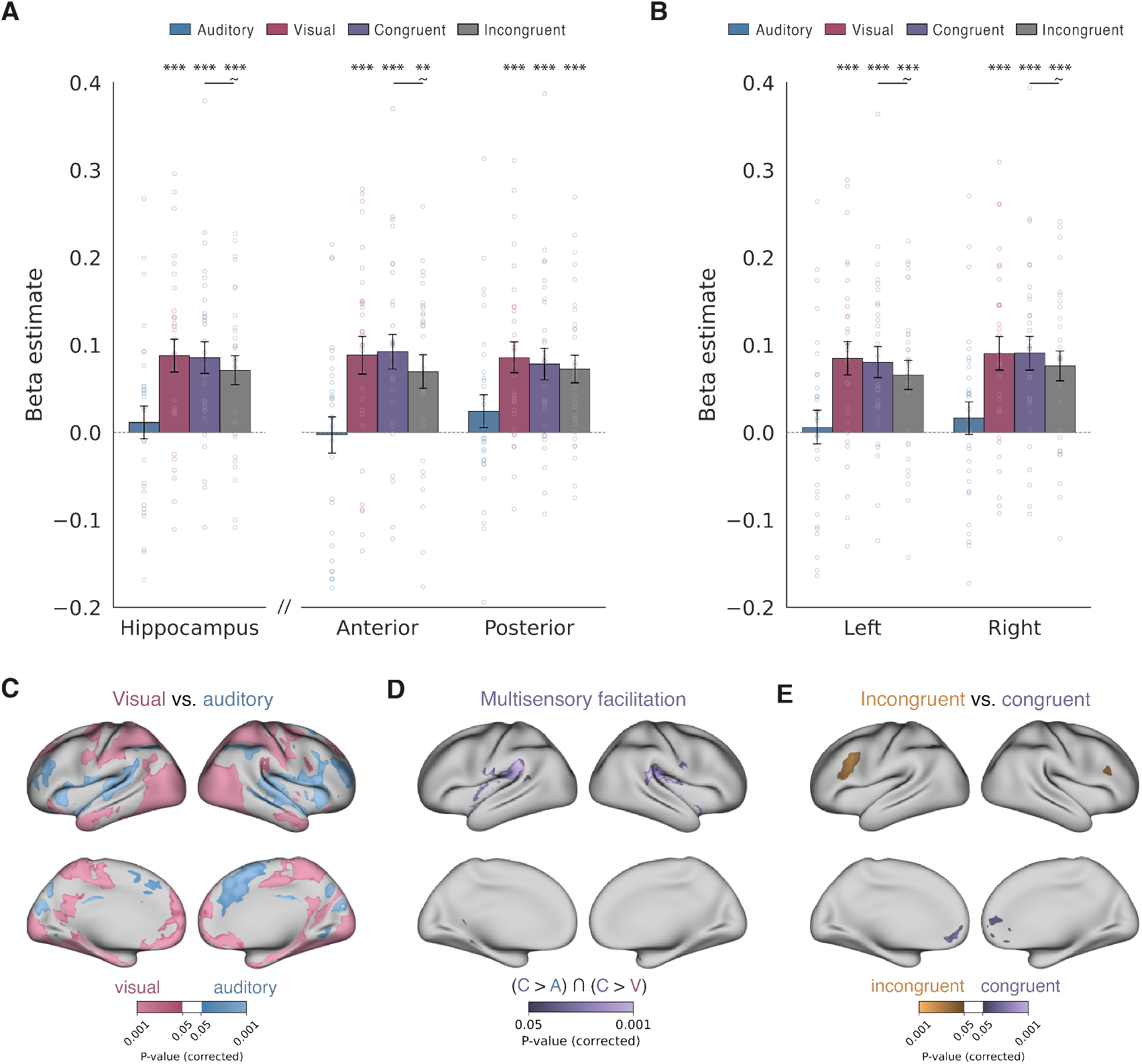
Univariate activation across the hippocampus and whole brain. **(A)** Mean beta estimates for each condition in the whole hippocampus and anterior/posterior subdivisions. **(B)** Mean beta estimates for each condition in the left and right hippocampus. **(C)** Group-level whole-brain auditory-only versus visual-only contrast. **(D)** Conjunction analysis identifying regions in which congruent audiovisual activation exceeded both auditory-only and visual-only conditions. **(E)** Whole-brain contrast of congruent versus incongruent audiovisual conditions. Individual participant data shown as circles. Error bars indicate SEM. ∼ *P <* 0.1, ^∗^*P <* 0.05, ^∗∗^*P <* 0.01, ^∗∗∗^*P <* 0.001. Whole-brain maps were TFCE corrected at *P <* 0.05.

Next, we estimated group-level effects in the whole brain using voxelwise non-parametric randomization tests (TFCE corrected at *P <* 0.05). A contrast of unisensory auditory versus visual conditions showed canonical activations in auditory and visual cortices, respectively (Figure 2C). To assess multisensory facilitation, we performed a conjunctive analysis of voxels in which activation for the congruent audiovisual condition significantly exceeded both of the unisensory auditory and visual conditions. This contrast revealed activity in several regions, including bilateral parietal operculum, insula, angular gyrus, temporal pole, and subcortical activity in the thalamus (Figure 2D). Finally, a contrast between congruent and incongruent audiovisual conditions revealed greater activation in bilateral medial prefrontal cortex for congruent and in left lateral prefrontal cortex for incongruent (Figure 2E). Together, these whole-brain effects validate the paradigm by replicating established patterns of unisensory and multisensory processing in the neocortex. The full list of peak coordinates for these contrasts is provided in Table S2. Critically, while these univariate results confirm that the hippocampus is sensitive to the visual modality, they provide no evidence of sensitivity to the auditory modality or for multisensory facilitation.

### Multivariate representations

We estimated single-trial activity patterns using GLMSingle ([29]) and performed RSA with a leave-one-run-out cross-validation approach. For a given ROI, we extracted voxel-wise beta coefficients for each movie of a condition in every run (Figure 3A). On each fold, we correlated the patterns from one held-out run with the average patterns from the other runs (Figure 3B), yielding a 10 (movie) by 10 (movie) similarity matrix for each condition. All correlation coefficients were Fisher z-transformed and the similarity matrices were averaged across folds. Diagonal elements corresponded to within-movie similarity (same movie across runs), while off-diagonal elements reflected between-movie similarity (different movies across runs). To quantify representational distinctiveness for each condition, we computed movie-specific pattern similarity (MSPS) as the signed difference between average within-movie and between-movie similarities (mean of diagonal cells minus mean of off-diagonal cells), with positive values indicating more distinctive neural representations for individual movie clips. MSPS was computed for each participant and statistical reliability for auditory, visual, and congruent conditions was calculated against zero (null hypothesis of no diagonal minus off-diagonal; i.e., that pattern similarity was not movie-specific).

**Figure 3:**
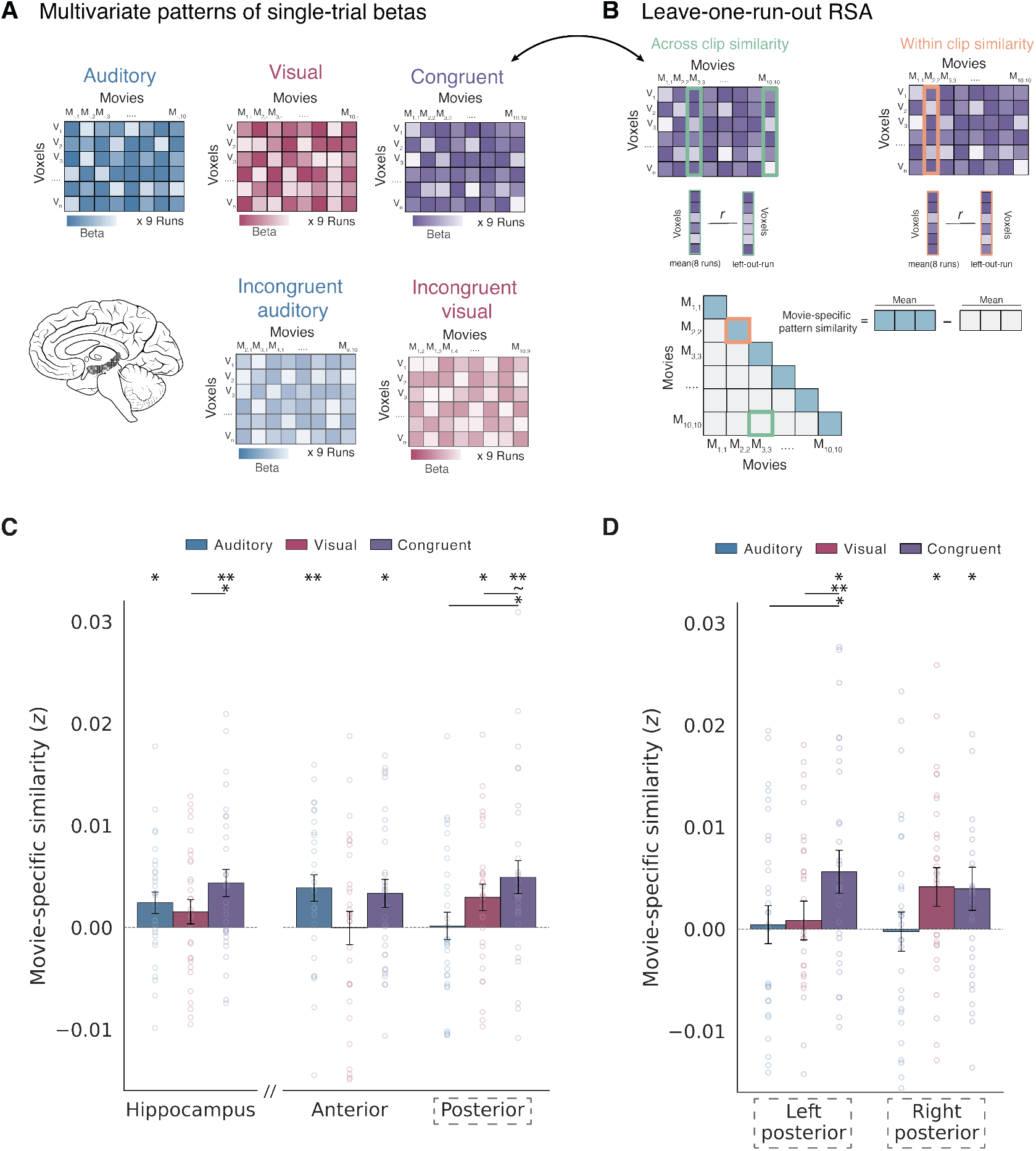
Movie-specific multivariate representations in the hippocampus. **(A)** To perform RSA, single-trial beta estimates were first extracted from a given ROI, generating multivariate activity patterns for each movie presentation. **(B)** We computed cross-validated RSA matrices independently for each condition using leave-one-run-out procedures (congruent audiovisual condition shown). Voxel-wise correlations were computed between movies (green) and within movies (orange) and averaged across cross-validation folds to construct the RSA matrix (bottom). To quantify movie-specific pattern similarity (MSPS), we computed the signed difference between mean diagonal (within-movie) and off-diagonal (between-movie) similarity values. **(C)** MSPS across conditions for the whole hippocampus (left) and anterior and posterior subdivisions (right). The posterior hippocampus showed enhanced pattern similarity for congruent audiovisual stimuli relative to unisensory conditions (grey hatched box). **(D)** MSPS across conditions for the left and right posterior hippocampus. Six data points outside the y-axis range are not shown; the range was chosen to match that of panel C to ease comparisons. Individual participant data shown as circles. Error bars indicate SEM. ∼ *P <* 0.1, ^∗^*P <* 0.05, ^∗∗^*P <* 0.01, ^∗∗∗^*P <* 0.001.

We found reliable MSPS for auditory, visual, and congruent movie clips in a subset of hippocampal ROIs (Figure 3C). Unisensory auditory representations were observed in the whole hippocampus (*P* = 0.017) and anterior hippocampus (*P* = 0.007). Unisensory visual representations were observed in the posterior hippocampus (*P* = 0.031). Congruent audiovisual representations were found in the whole hippocampus (*P* = 0.001), anterior hippocampus (*P* = 0.012), and posterior hippocampus (*P* = 0.003). The subiculum (*P* = 0.031) and DG (*P* = 0.031) showed significant congruent audiovisual MSPS but no significant unisensory MSPS (all *P >* 0.05; Figure S1C). In MTL cortex (Figure S3B), PHC showed significant MSPS for the visual (*P <* 0.001), auditory (*P* = 0.002), and congruent audiovisual (*P <* 0.001) conditions, and PRC showed significant MSPS only for the congruent audiovisual MSPS (*P* = 0.027); EC showed no significant effects (all *P >* 0.05.)

The posterior hippocampus showed evidence of multisensory facilitation, with MSPS for the congruent condition significantly greater than for the auditory condition (*P* = 0.013) and marginally greater than for the visual condition (*P* = 0.084). The whole hippocampus showed a similar pattern, with enhanced MSPS for the congruent condition relative to both the visual (*P* = 0.029) and auditory conditions, though the latter did not reach significance (*P* = 0.15). We did not observe a facilitation pattern in the anterior hippocampus or in any hippocampal subfield (Figure S1C). Examining hemispheres separately (Figure 3D), multisensory facilitation was lateralized to the left posterior hippocampus, where congruent MSPS was significant (*P* = 0.011) and exceeded both the auditory (*P* = 0.040) and visual (*P* = 0.009) conditions. In contrast, the right posterior hippocampus showed significant visual-only (*P* = 0.039) and congruent audiovisual (*P* = 0.036) MSPS, but no facilitation. Lateralization in the whole and anterior hippocampus is shown in Figure S4A–B; full descriptive statistics for all ROIs are provided in Tables S3 and S4.

Critically, congruent audiovisual MSPS could reflect a conjunctive audiovisual code, in which congruent stimuli evoke movie-specific structure not present in either modality alone, or simply a linear combination of the unisensory representations. To test this, we ran a cross-validated multiple regression, modeling the congruent audiovisual similarity matrix as a function of the auditory-only and visual-only similarity matrices (Figure 4A). Specifically, we estimated these coefficients by predicting each held-out congruent run from the remaining unisensory runs, following our main RSA approach (see Methods). Averaging across folds, we derived the auditory and visual regression coefficients, the variance explained (*R*^2^), and the residual congruent audiovisual MSPS. The visual-only coefficient was significant in the whole hippocampus (*P* = 0.003) and posterior hippocampus (*P* = 0.005; Figure 4B), whereas the auditory-only coefficient was not significant in any hippocampal ROI (all *P >* 0.05). This visual dominance was expected, as the regression models the full similarity matrix rather than isolating movie-specific structure. Critically, the unisensory matrices explained no significant variance in any hippocampal subdivision (all *P >* 0.05). This was in stark contrast to visual and auditory sensory ROIs in cortex, which accounted for substantial variance (*R*^2^ ≈ 11–36%; for full results in sensory ROIs, see Figure S5). Finally, the residual congruent audiovisual MSPS remained above baseline in the whole hippocampus (*P* = 0.002), anterior hippocampus (*P* = 0.019), and posterior hippocampus (*P* = 0.005; Figure 4C). Together, these findings indicate that the hippocampus carries movie-specific audiovisual structure that is not reducible to its unisensory components. For full statistical reporting, see Table S5

**Figure 4:**
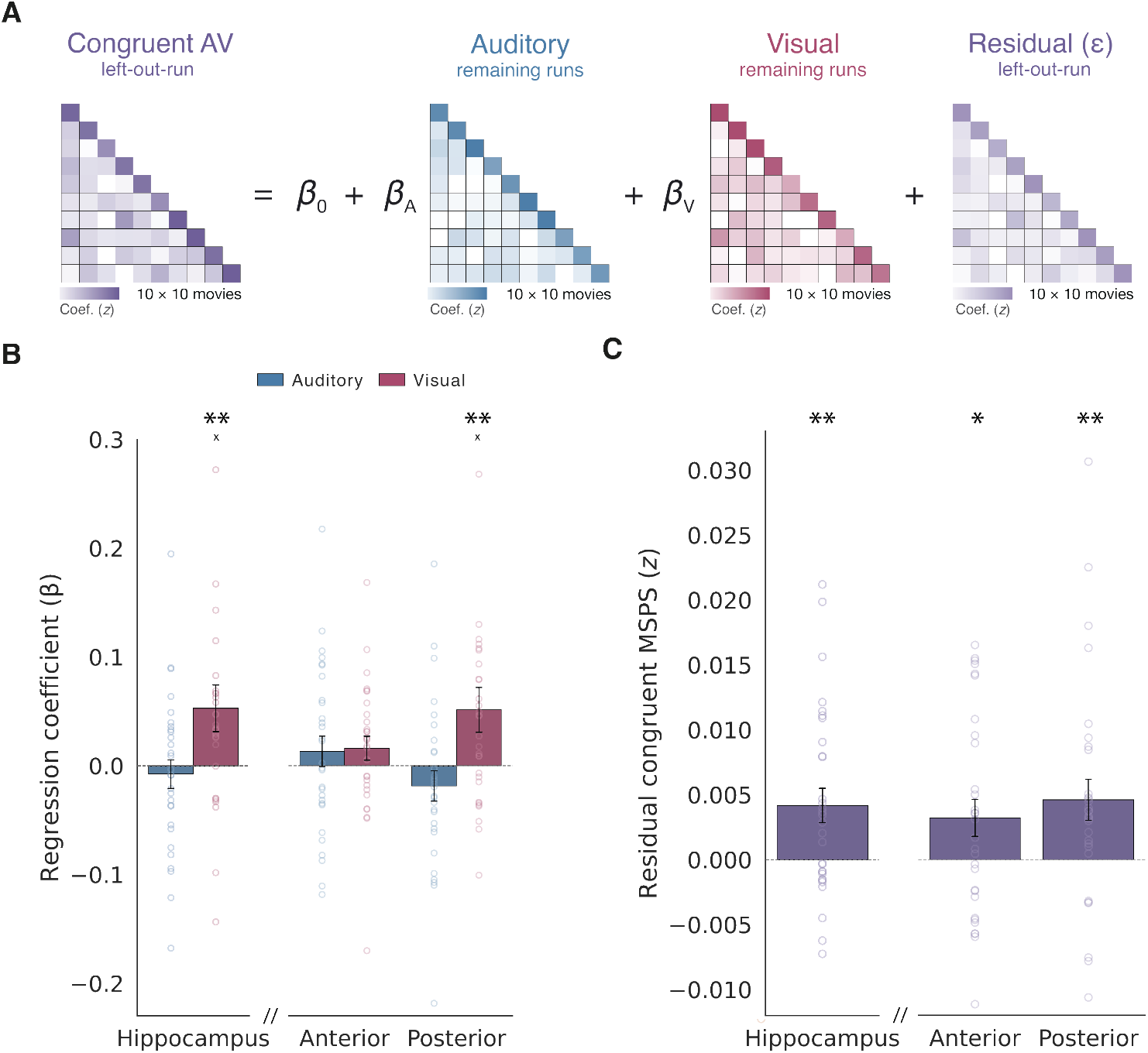
Conjunctive audiovisual coding in the hippocampus. **(A)** To test whether congruent audiovisual representations could be explained by a linear combination of unisensory representations, we performed a cross-validated multiple regression modeling the congruent audiovisual similarity matrix as a function of the auditory-only and visual-only similarity matrices. Regression coefficients (*β_A_*, *β_V_*) were estimated by iteratively predicting the congruent matrix on a held-out run from the remaining runs, with estimates averaged across folds. **(B)** Auditory and visual regression coefficients are shown for the whole hippocampus and anterior/posterior subdivisions. **(C)** Residual congruent audiovisual MSPS after regressing out the auditory-only and visual-only contributions. Individual participant data shown as circles; error bars indicate SEM. × markers indicate individual data points outside the plotted y-axis range. ^∗^*P <* 0.05, ^∗∗^*P <* 0.01

Note that this multivariate analysis could not be performed straightforwardly in the incongruent condition because each run contained a unique audiovisual combination. However, each modality still carries movie-specific information across runs; for example, an ambulance audio track is always paired with a distinct visual component within each run. This results in “incongruent visual” and “incongruent auditory” matrices, allowing us to test whether incongruence suppresses pattern similarity relative to the unisensory conditions (despite both having stable audio or video). Although we did not find any significant differences in the hippocampus (Figure S6A), we observed a significant suppression of MSPS for the incongruent relative to unisensory conditions in the PHC (Figure S6B) and at the whole brain level (Figure S7).

To test whether MSPS was modality-specific or amodal, we cross-correlated the voxelwise patterns for auditory and visual presentations of the same versus different movies using the cross-validated leave-one-run-out RSA approach (Figure 5A). Reliable MSPS in this analysis would indicate the existence of movie-specific representations invariant to sensory modality. The whole hippocampus (*P* = 0.003) and the anterior hippocampus (*P* = 0.003) exhibited significant crossmodal transfer (Figure 5B), whereas the posterior hippocampus did not (*P* = 0.22). Crossmodal transfer was significant in the right anterior hippocampus (*P* = 0.016) but not the left (*P* = 0.20; Figure 5C), with the same asymmetry at the level of the whole hippocampus (right: *P* = 0.006; left: *P* = 0.26; Figure S8A). The direct contrast between hemispheres was not significant in either ROI (both *P >* 0.05). In hippocampal subfields (Figure S9A), there was marginal evidence of crossmodal transfer in DG (*P* = 0.091) and subiculum (*P* = 0.091). In MTL cortex, PHC showed robust crossmodal transfer (*P <* 0.001; Figure S9B). See full descriptive statistics in Table S4.

**Figure 5:**
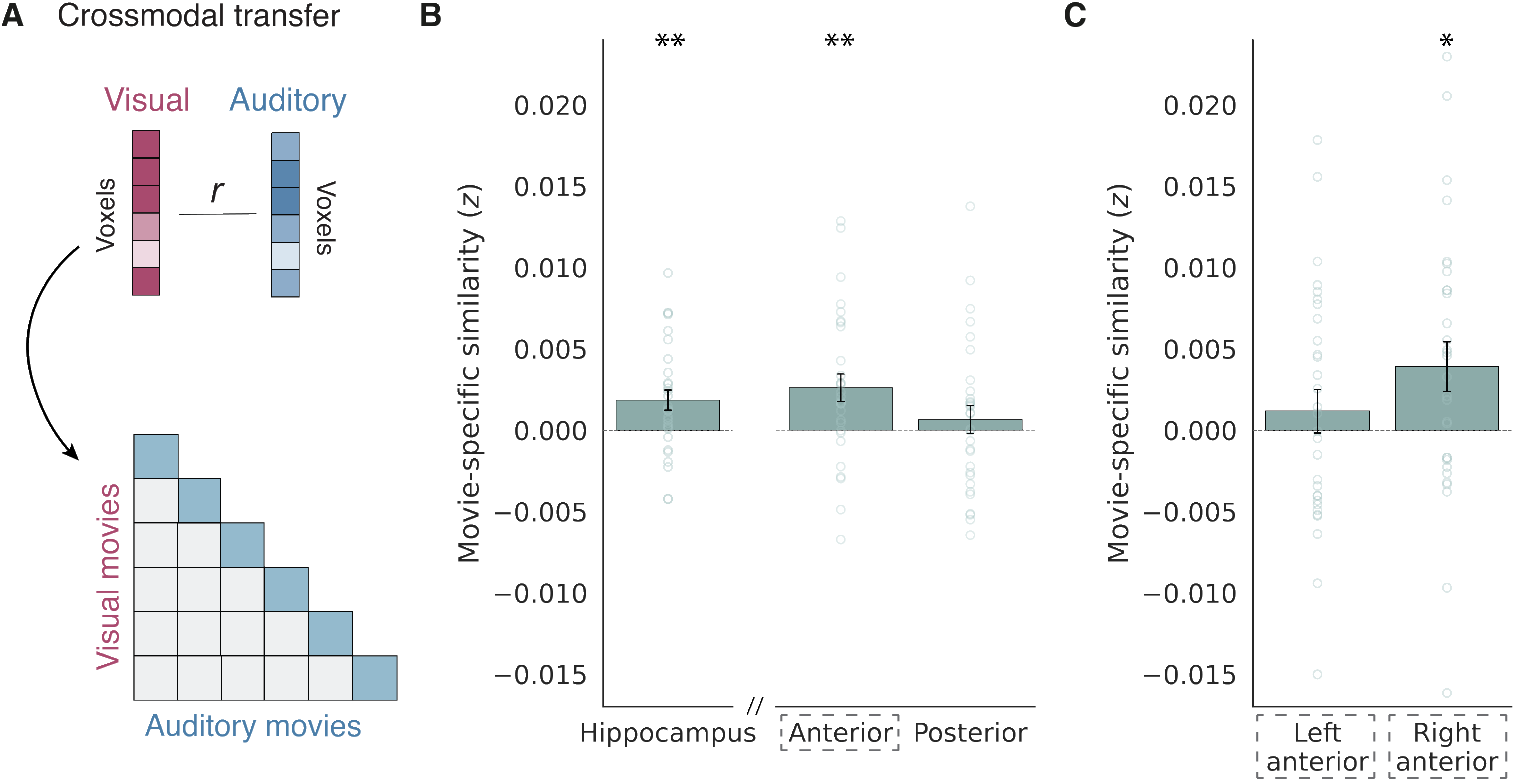
Crossmodal transfer in the hippocampus. **(A)** To estimate crossmodal transfer, we extended the cross-validated RSA approach by computing voxel-wise correlations between auditory-only and visual-only scene presentations. Movie-specific pattern similarity across modalities was quantified as the signed difference between same-movie (diagonal) and different-movie (off-diagonal) correlations. **(B)** Crossmodal MSPS in the whole hippocampus and anterior/posterior subdivisions. **(C)** Crossmodal MSPS for the left and right anterior hippocampus. Individual participant data shown as circles; error bars indicate SEM. ^∗^*P <* 0.05, ^∗∗^*P <* 0.01.

The multivariate effects reported above could in principle benefit from memory retrieval, with repeated stimulus exposure allowing for reinstatement of movie-specific details or mental imagery of the absent modality. If so, we would expect that the magnitude of these effects should increase across repeated stimulus presentations. We tested this possibility in two complementary analyses. First, we leveraged the leave-one-run-out cross-validation RSA procedure, in which each run is held out and correlated with the average of the remaining runs. This yields 9 estimates (one per fold), each corresponding to a run’s position in the task. We therefore labeled the cross-validated folds by held-out run position (runs 1–9) and correlated run position with the magnitude of our effects of interest (multisensory facilitation and crossmodal transfer) in that run. If repeated exposure strengthened retrieval- or imagery-based contributions, we would expect a positive correlation between run position and effect magnitude. Neither multisensory facilitation in the posterior hippocampus (congruent audiovisual *>* visual MSPS; Spearman’s *ρ* = 0.055, *P* = 0.38) nor crossmodal transfer in the anterior hippocampus (*ρ* = 0.037, *P* = 0.55) strengthened with more stimulus repetition across runs. Second, we ran a split-half analysis comparing effects estimated within the first four runs versus the last four runs (omitting the middle run for balance; Figure S10). In the first half, the posterior hippocampus showed significant MSPS for the auditory-only (*P* = 0.017) and congruent audiovisual (*P* = 0.050) conditions; in the second half, there were no significant effects. In contrast, crossmodal transfer in the anterior hippocampus was significant in the second half (*P* = 0.005) but not the first half (*P* = 0.12). However, neither multisensory facilitation nor crossmodal transfer differed significantly between the two halves (both *P >* 0.05). These results raise the possibility that facilitation and transfer might be affected differently by stimulus repetitions, perhaps because crossmodal transfer involves learning amodal abstractions.

To examine multivariate results beyond the hippocampus, we conducted an exploratory whole-brain searchlight analysis. In each participant, we applied the cross-validated leave-one-run-out RSA analysis within a cubic searchlight centered on each brain voxel and assigned the resulting MSPS value to that voxel (Figure 6A). We then assessed reliability across participants (TFCE corrected at *P <* 0.05). A contrast between unisensory auditory and visual conditions showed robust MSPS in auditory and visual cortices, respectively (Figure 6B). Next, we performed a conjunctive contrast to identify regions in which congruent audiovisual stimuli exceeded both unisensory conditions. This revealed multisensory facilitation across the superior temporal lobe, including posterior STG and adjacent posterior perisylvian cortex (planum temporale and parietal operculum), as well as in the insula, temporal pole, and amygdala (Figure 6C). Crossmodal transfer effects at the whole-brain level were robust across perisylvian cortex, angular gyrus, precuneus, and lateral visual cortex (Figure 6D). The full list of peak coordinates for these contrasts is provided in Table S6.

**Figure 6:**
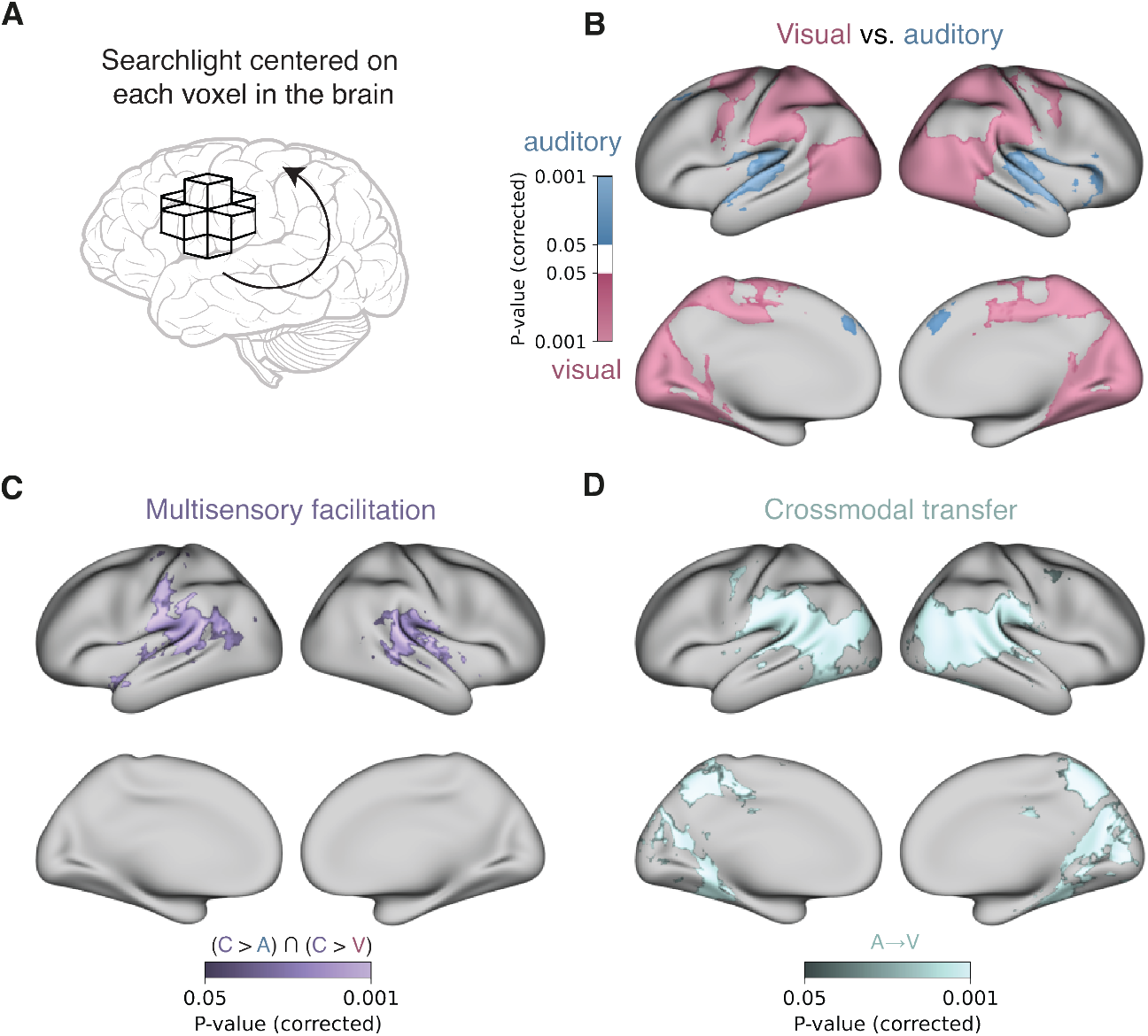
Whole-brain searchlight RSA results. **(A)** An exploratory searchlight was applied to estimate multivariate RSA effects in regions across the whole brain **(B)** Group-level auditory-only versus visual-only MSPS contrast. **(C)** Conjunctive map showing regions in which congruent audiovisual MSPS exceeded both auditory-only and visual-only conditions. **(D)** Above-baseline crossmodal transfer MSPS. All maps were TFCE corrected at *P <* 0.05.

## Discussion

This study focused on the representation of unisensory and multisensory content in the human hippocampus. We found evidence of auditory and visual coding, multisensory facilitation, and crossmodal transfer along the hippocampal long axis. Namely, audiovisual clips were represented more reliably than auditory and visual versions of the same clips in the posterior hippocampus, whereas these unisensory versions shared an abstract representation in the anterior hippocampus. These findings are consistent with a posterior-to-anterior gradient in the hippocampus from fine-grained to abstract representations, respectively [26, 27]. Multisensory processing in the hippocampus was also lateralized, with clearer facilitation in the left (posterior) hippocampus and clearer crossmodal transfer in the right (anterior) hippocampus; this is another case of hemispheric specialization in the human hippocampus [30].

### Auditory scene representation in the medial temporal lobe

We observed a clear distinction between analytical approaches: univariate analyses detected only visual responses across the hippocampus, with no activation for auditory stimuli, whereas multivariate pattern analyses revealed representations of auditory scenes. Specifically, there was significant movie-specific pattern similarity for auditory-only stimuli in the whole hippocampus and anterior hippocampus. This suggests that auditory content can be encoded in distributed voxel-wise patterns and not readily captured in the mean activity across these voxels, underscoring the need for multivariate approaches to detect non-visual representations in the human hippocampus.

This dissociation may be theoretically informative, as opposed to simply reflecting the greater sensitivity of multivariate methods. Whereas the univariate analysis reflects overall regional engagement, our multivariate RSA indexes stimulus-specific information carried across distributed voxel patterns. The movie-specific (event-level) structure captured in this analysis is consistent with prominent accounts implicating the hippocampus in the rapid, sparse coding of individuated events [31, 32, 33]. Importantly, this does not preclude a univariate hippocampal response to auditory stimuli, which would be expected given extensive evidence for auditory processing in the hippocampus, primarily observed in non-human animals [5]. Univariate auditory responses may be identified under conditions that more strongly drive hippocampal activity, such as paradigms imposing explicit memory demands or those not relying on stimulus repetitions.

Auditory and visual stimuli may differ in perceptual salience or attentional demands, complicating the comparison between unisensory conditions. Several aspects of our data argue against a salience-driven account. First, response times in the offset detection behavioral task did not differ between unisensory conditions, indicating relatively matched task engagement (*P >* 0.05). Second, the incongruent audiovisual condition elicited lower univariate responses than both unisensory conditions across hippocampal ROIs (Figure 2A–B); if low-level salience in either modality were driving these responses, adding a second discordant modality should have enhanced them. Third, although salience could in principle modulate the overall magnitude of BOLD responses, our central effects are defined by movie-specific pattern similarity, which controls for the pattern similarity across different movies; as such, our multivariate analyses are inherently robust to condition-level differences in salience or attention.

Prior evidence for auditory coding in the human MTL is limited (cf. [34, 35]). Yet, an extensive body of work has established a role for the hippocampus in visual scene perception [10, 11]. Our findings provide initial evidence that the hippocampus may additionally support auditory scene perception in the absence of explicit memory demands, consistent with accounts of *domain-general* relational coding in the human hippocampus [36, 37]. Testing this hypothesis more directly will require establishing a preference in the hippocampus for auditory scenes over other forms of auditory stimuli, such as auditory objects or simple acoustic stimuli, extending experiments from the visual modality [16, 38]. It would further be interesting to test whether the hippocampus is specifically sensitive to the global structure or local details of auditory scenes, as has been examined in the visual modality [17].

We additionally observed robust auditory coding in PHC — a major input structure to the hippocampus via the entorhinal cortex. Although traditionally considered a visual region involved in place and scene processing, our findings align with recent evidence that the PHC responds to spoken descriptions of spatial information and landmarks [34]. Importantly, these effects may reflect crossmodal interactions, as evidenced by stronger visual effects and robust crossmodal generalization in our data. Nevertheless, these findings suggest a potential role for PHC in representing auditory “landscapes” that parallels its established function in visual scene processing.

### Multisensory facilitation in the posterior hippocampus

We found enhanced movie-specific pattern similarity in the posterior hippocampus for congruent audiovisual stimuli relative to both unisensory modalities, especially in the left hemisphere. Incongruent audiovisual stimuli did not show enhanced pattern similarity relative to unisensory conditions, further supporting the sensitivity of posterior hippocampus to multisensory congruence. We did not observe multisensory facilitation in the anterior hippocampus or in any individual hippocampal subfields. Together, we interpret these findings as facilitation of perceptual processing in the posterior hippocampus when a matching and synchronized stimulus from another modality is present. This aligns with prior evidence that the posterior hippocampus exhibits stronger connectivity with sensory cortices and preferential coding of fine-grained visual features [39, 26, 40].

Importantly, these congruent audiovisual representations were not reducible to their unisensory components. Using a cross-validated multiple regression, we found that movie-specific pattern similarity for congruent stimuli remained reliably above zero after regressing out the auditory-only and visual-only similarity matrices. This indicates a conjunctive audiovisual code in the hippocampus, in which congruent stimuli elicited unique movie-specific structure. This contrasts with sensory cortical ROIs, in which the unisensory matrices accounted for substantial variance in the congruent representation. Nevertheless, residual conjunctive structure was evident in both sensory areas and the hippocampus.

A whole-brain searchlight contrast between congruent audiovisual and unisensory conditions revealed robust effects in the STS and posterior STG. This aligns with an extensive literature implicating these regions in audiovisual integration (e.g., [22, 23, 24, 25]). These findings situate the posterior hippocampus within a broader network supporting multisensory processing, raising questions about the distinct functional roles of these regions. For instance, does audiovisual facilitation have different consequences for multisensory perception versus memory across these sites? Furthermore, the functional interaction between the hippocampus and these cortical sites represents an important direction for future research.

### Crossmodal generalization in the anterior hippocampus

Studies of multisensory perception interpret crossmodal transfer as reflecting “equivalence representations,” reflecting abstracted, modality-invariant contents [41, 42, 43, 44]. The anterior (but not posterior) hippocampus exhibited crossmodal transfer, such that multivariate patterns belonging to the same scene’s unisensory auditory and visual stimuli were more similar to each other than to patterns evoked by unisensory stimuli from different scenes. In the visual modality, the anterior aspects of the hippocampus have been implicated in representing abstract associations as opposed to perceptual details [39, 27]. Notably, recent work has found that neighboring anterior temporal lobe structures show similar crossmodal object coding, with experience attenuating a visual bias in perirhinal cortex [35]. Our whole-brain searchlight analysis found crossmodal transfer in the STS, pSTG, as well as high-level visual areas, in line with research into amodal representations across vision and audition in cortical sites [45, 46].

It is important to acknowledge that such effects can emerge from a range of shared contents across sensory modalities [41], including shared semantic content, low-level temporal coherence (e.g., energy fluctuations), or reinstatement of the absent modality via pattern completion or imagery. The present movie-based paradigm is not well-suited for adjudicating between these interpretations, but other paradigms that systematically manipulate stimulus features could be informative in future studies. Although our analyses across repeated exposures provided limited support for a retrieval-based account of these effects, we observed a trend toward enhanced crossmodal transfer in later runs, which suggests that these effects may be driven in part by learned audiovisual associations (Figure S10). In fact, even without an increase across runs, it is possible that the pre-experimental exposure phase enabled retrieval processes even early in the experiment. Consistent with this possibility, we observed evidence of perceptual reinstatement in both modalities: auditory-only MSPS was significant in visual cortical ROIs, and visual-only MSPS was significant in auditory cortical ROIs (Figure S3). These multivariate effects were notably weaker than the patterns for the presented modality. Even so, they suggest some degree of reinstatement of absent modalities. Future work that directly examines crossmodal learning will be necessary to fully adjudicate between perceptual- and retrieval-based accounts.

### Implications for hippocampal long-axis specialization and lateralization

We interpret our findings in terms of a posterior-to-anterior gradient from fine-grained to abstract representations, which provides a compelling account of the observed multisensory effects [26, 27]. Nevertheless, other accounts of long-axis specialization offer alternative interpretations. One prominent account distinguishes anterior encoding from posterior retrieval [47], raising the possibility that posterior audiovisual facilitation reflects enhanced recall of congruent stimuli. A more recent network-driven account instead links the posterior hippocampus to salience and target detection, and the anterior hippocampus to memory and scene construction [48]. This is broadly consistent with our analyses across repeated exposures, which showed a trend toward stronger crossmodal transfer in the anterior hippocampus (Figure S10). Adjudicating between these accounts will require further work, particularly designs that dissociate memory from perceptual and representational contributions to hippocampal function.

Our results additionally revealed hemispheric asymmetries along the long axis: multisensory facilitation was strongest in the left posterior hippocampus, whereas crossmodal transfer was driven by the right anterior hippocampus. The human hippocampus displays strong lateralization across a range of memory and navigation tasks [30, 49, 50, 51, 52, 53]. Our results are broadly consistent with a left-hemisphere bias for episodic coding of stimulus content and a right-hemisphere bias for coarser, configural representations that are better suited to abstracting across modality-specific detail. The latter aligns with work showing a right hippocampal bias for statistical learning across different types of visual stimuli [54, 55]. These interpretations are speculative, as the principles governing hippocampal laterality remain poorly understood. Even so, the asymmetries observed here motivate the use of multisensory stimuli to constrain hypotheses regarding lateralization in the human hippocampus.

### Conclusions

Despite decades of research on the human hippocampus, we lack a basic understanding of how this brain structure processes non-visual perceptual information and integrates across senses. By providing a comprehensive initial account of how multisensory coding is organized in the human hippocampus, this work provides a foundation for bridging across perception, memory, language, and other domains. A critical next step will be to link these unisensory and multisensory representations to the formation of episodic memories.

## Methods

### Participants

Thirty-two young adults with normal or corrected-to-normal vision were recruited from Yale University and the New Haven communities. Two participants were excluded due to excessive head motion. The remaining participants comprised our final sample (N = 30; 16 female; mean age = 24.78 years, age range = 18.80–37.53). Although the effect sizes were unknown preventing a power analysis, this sample size is consistent with, or greater than, other studies of hippocampal function [56]. All participants provided written informed consent and were compensated for their time ($25 per hour). The study was approved by the Yale University Institutional Review Board (protocol number: 2000022976).

### Stimuli generation

To generate the stimuli, we sourced 12 naturalistic audiovisual movies. Nine movies were from YouTube and three were from Pexels (https://www.pexels.com). In searching for stimuli, we prioritized movies that varied in content (animate/inanimate, indoor/outdoor, etc.) and that maximized audiovisual coherence (i.e., the visual and auditory components were clearly aligned to specific events). Only movies that contained high-fidelity video (1080p or 4K) and audio (48 kHz) were included. Movies containing legible speech were not considered. The final 12 movies included a ping-pong game, wolves howling, waves breaking, birds chirping, race cars passing by, and other multisensory scenes (Figure 1A).

Using Premiere Pro 2024 (Adobe Inc.), we cropped each movie into a 6-s segment. These segments were loudness normalized to the EBU R128 loudness standard with a target loudness of −23 LUFS. The movies were exported at 1280 x 720 resolution at 30 frames per second, with H.264 video codec and AAC audio codec. The audio was configured as stereo across all movies at 24 kHz.

From these processed movies, we selected two movies to serve as stimuli for the practice block, while the remaining 10 movies made up the experimental stimuli. To construct the task conditions, we used Premiere Pro to modify the movies. For the visual-only condition, we extracted the video component while removing the audio. For the auditory-only condition, we extracted the audio component while removing the video (Figure 1B). To create the incongruent condition, we re-paired the audio and visual components from different movies. This resulted in 90 possible incongruent stimuli (10 videos × 9 non-matching audio tracks each). For the practice block, we created two incongruent stimuli using the same mismatching procedure. In total, the experimental stimulus set comprised 120 stimuli: 10 congruent audiovisual movies (original movies), 10 visual-only, 10 auditory-only, and 90 incongruent audiovisual movies. The practice block contained a total of 6 stimuli. Source URLs for all movies and the specific time segments extracted are available on the OSF project (https://osf.io/jcg8a).

### Task design and procedures

The experiment consisted of 9 experimental runs, each containing 40 movie presentations: 10 congruent audiovisual, 10 auditory-only, 10 visual-only, and 10 incongruent audiovisual. All congruent, auditory-only, and visual-only stimuli were presented in every run. For the incongruent condition, each run contained all 10 video components and all 10 audio components, each paired with a mismatched audio or visual track. That is, each video was paired with audio from a different movie, with pairings varying across runs (e.g., ambulance video paired with ping-pong audio in one run, with waves audio in another, and so on). This ensured that the visual and auditory content from all movies appeared with equal frequency in every run while varying the specific incongruent pairings across runs, enabling leave-one-run-out cross-validation comparable to the other conditions. The order of stimuli in each run was pseudo-randomized by shuffling trials in each half of the run, ensuring an equal number of stimuli from all conditions in both halves. Finally, the order of the runs themselves was randomized across participants, to prevent order effects for the incongruent condition (the runs were all identical for the congruent, auditory-only, and visual-only conditions). In each run, the movies were shown sequentially with a jittered ITI (4.5, 6, and 7.5 s; Figure 1B). A central fixation cross was presented for the full 6 s of every trial, including during auditory-only trials, and was superimposed over videos in the visual-only and audiovisual conditions. The onset of each movie was triggered by the start of a brain volume acquisition to ensure that any timing delays were accounted for and did not accumulate across the experimental session.

Participants performed a simple cover task while viewing the movies to ensure attention. They were instructed to press a button with their index finger within 1 s of each movie’s offset (across all conditions). Visual feedback was provided after each trial, with a red fixation cross indicating an incorrect response (button press during the movie or >1 s after offset), while a green fixation indicated a correct response. At the end of each run, participants were shown their cover task accuracy (% correct responses). This paradigm was designed to maintain participants’ attention throughout each movie while imposing similar task demands across conditions. Participants made all responses on an MRI-compatible button box.

In the scanner, and prior to the experimental runs, participants passively viewed the 10 movies in their original audiovisual form to gain familiarity with the auditory and visual components. This pre-exposure intended to reduce memory demands during the main experiment, consistent with prior work [11]. Participants then completed one practice run that mirrored the experimental procedures but used the two held-out practice stimuli presented in each format.

### Audio delivery procedures

To deliver high-fidelity audio, we applied noise attenuation methods to overcome scanner noise. We used OptoActive inflatable headphones with active noise cancellation (OptoAcoustics, Mazor, Israel; https://www.optoacoustics.com), providing approximately 30 dB of noise suppression. Prior to the experimental session, we calibrated the active noise cancellation system based on the specific noise from the fMRI sequence while the participant was positioned in the scanner. Participants also wore semi-open earplugs, intended to provide an additional 10 dB of passive noise attenuation, while preserving stimulus transmission from the headphones. To maintain the spectral fidelity of auditory stimuli during active noise cancellation, we applied frequency-dependent equalization through the OptoAcoustics system. Following active noise cancellation calibration, we adjusted the volume level based on participants’ feedback when listening to an audiovisual clip not included in the experiment. This volume level was held constant throughout the experimental session.

### MRI acquisition

Data were acquired on a Siemens Prisma 3T scanner with a 64-channel volume head coil at BrainWorks in the Wu Tsai Institute at Yale University. Functional scans were collected using an echo-planar imaging (EPI) sequence with the following parameters: repetition time (TR) = 1,500 ms; echo time (TE) = 32 ms; 90 axial slices; voxel size = 1.5 × 1.5 × 1.5 mm; flip angle = 64°; multiband factor = 6. We also collected a pair of spin-echo scans with opposite phase-encoding directions for distortion correction (TR = 11,390 ms; TE = 66 ms). We acquired two anatomical scans: a high-resolution 3D T1-weighted (T1w) MPRAGE sequence (TR = 2,400 ms; TE = 2.19 ms; voxel size = 1 × 1 × 1 mm; 208 sagittal slices; flip angle = 8°), and a T2-weighted (T2w) turbo spin-echo sequence (TR = 11,170 ms; TE = 93 ms; 54 coronal slices; voxel size = 0.44 × 0.44 × 1.5 mm; distance factor = 20%; flip angle = 150°) positioned perpendicular to the anterior-posterior axis of the hippocampus to assist with automated segmentation of medial temporal lobe (MTL) regions of interest (ROIs). These sequence parameters have been used in recent studies of the hippocampus [57, 56].

### fMRI preprocessing

fMRI data were preprocessed using fMRIPrep version 23.2.1 [58], based on Nipype 1.8.6 [59]. The T1-weighted anatomical image was corrected for intensity non-uniformity using N4BiasFieldCorrection [60] distributed with ANTs 2.5.0 [61]. The corrected T1w image was skull-stripped using an implementation of the antsBrainExtraction.sh workflow. Brain tissue segmentation of cerebrospinal fluid (CSF), white matter (WM), and gray matter (GM) was performed on the brain-extracted T1w image using FAST in FSL ([62]). Brain surfaces were reconstructed using recon-all from FreeSurfer 7.3.2 [63]. Volume-based spatial normalization to standard space was performed through nonlinear registration with antsRegistration (ANTs 2.5.0) to the MNI152 standard template [64].

For each of the nine functional runs, slice-timing correction was applied. Head motion was corrected using mcflirt in FSL ([65]), and head-motion parameters (six rotation and translation parameters) were estimated for use as nuisance regressors. Susceptibility distortion correction was performed using a fieldmap-based approach, with field inhomogeneities estimated from spin-echo images with opposite phase-encoding directions using topup in FSL [66]. Fieldmaps were not acquired correctly for the first three participants and so susceptibility distortion correction was not applied in these cases. The corrected BOLD reference image was co-registered to the T1w reference using bbregister in FreeSurfer, which implements boundary-based registration [67] with six degrees of freedom.

We excluded trials in which framewise displacement exceeded 1.5 mm on any TR. Runs with more than two excluded trials were removed entirely, and participants with more than two excluded runs were removed from the dataset. This procedure resulted in the exclusion of two participants. In the remaining sample (N = 30), one run was excluded from each of two participants and 20 individual trials total were excluded across all participants.

### Defining regions of interest

MTL structures were segmented from each participant’s T1-weighted and T2-weighted anatomical images using the Automatic Segmentation of Hippocampal Subfields (ASHS; [68]) machine learning toolbox. The ASHS algorithm was trained on 24 manual segmentations of hippocampal subfields [69, 70]. This procedure generated participant-specific masks for hippocampal subfields CA1, CA2/3, and dentate gyrus (DG), as well as extrahippocampal MTL regions including entorhinal cortex (EC), perirhinal cortex (PRC), and parahippocampal cortex (PHC). Anterior and posterior hippocampal ROIs were manually defined for each participant on their native-space T1w image as the most posterior coronal slice in which the uncus was visible [26, 28]. We defined masks for the occipital pole (OP), lateral occipital cortex (LOC), Heschl’s gyrus (HG), and posterior superior temporal gyrus (pSTG) from the Harvard-Oxford atlas [71].

### Univariate GLM analysis

We constructed event-related general linear models (GLMs) using Nilearn version 0.9.2 [72]. For each participant and run, we included four condition regressors: auditory-only, visual-only, congruent audiovisual, and incongruent audiovisual. Each trial was modeled as a 6-s boxcar function aligned to stimulus onset and convolved with a canonical hemodynamic response function (HRF) using the Glover model with temporal and dispersion derivatives [73]. Additional regressors included six rigid-body motion parameters from fMRIPrep and nuisance regressors derived from high-variance voxels and high-pass filtering. The GLM was fit using an autoregressive AR(1) noise model. No spatial smoothing was applied during model estimation.

For ROI analyses, we extracted subject-specific beta estimates from each ROI and averaged across voxels within region. For whole-brain analyses, subject-level contrast maps were transformed from native T1w space to MNI space using ANTs [61] and spatially smoothed with a 5 mm FWHM Gaussian kernel. Non-parametric randomization tests were applied using randomise in FSL ([74]).

### Representational similarity analyses

We estimated single-trial activity patterns using GLMSingle [29], which modeled each 6-s movie presentation and generated regularized beta estimates for each trial and voxel. GLMSingle applied voxel-wise HRF optimization. Furthermore, using GLMdenoise, this procedure automatically estimated nuisance regressors and executed noise correction [75]. Finally, regularization was performed to improve the final beta estimates.

For each ROI, we extracted single-trial beta estimates for each voxel, generating a multivariate activity pattern for each trial. For example, each of the 10 auditory-only stimuli was presented once per run across 9 experimental runs, yielding 90 beta patterns for the auditory-only condition (assuming no trial exclusions due to motion). Similarly, we obtained patterns each for the congruent audiovisual, visual-only, and incongruent audiovisual conditions (Figure 3A).

For each condition, we computed a cross-validated representational similarity matrix using leave-one-run-out procedures (Figure 3B). We iteratively left out one run and correlated it with the average of the remaining eight runs, repeating this procedure across all cross-validation folds. All correlation coefficients were Fisher z-transformed before averaging across folds. To quantify pattern distinctiveness for each condition and participant, we computed the difference between the mean diagonal z-value (within-scene similarity) and the mean off-diagonal z-value (between-scene similarity). Higher values indicate more distinct multivariate representations for individual movies.

To assess crossmodal transfer, we applied the same RSA procedure but compared patterns between the auditory-only and visual-only conditions (Figure 5A). Specifically, we correlated multivariate patterns for the same scene presented in different modalities (diagonal elements) and for different scenes across modalities (off-diagonal elements). As before, we computed the difference between mean within-scene similarity (diagonal z-values) and mean between-scene similarity (off-diagonal z-values) to quantify crossmodal transfer within individual scenes.

### Cross-validated multiple regression on similarity matrices

To assess whether the congruent audiovisual representations could be explained by a linear combination of unimodal sensory representations, we performed a cross-validated multiple regression in which the congruent audiovisual similarity matrix was modeled as a function of the auditory-only and visual-only similarity matrices. For each participant and ROI, we fit the following regression over the cells of the 10 (movie) by 10 (movie) Fisher z-transformed similarity matrices:

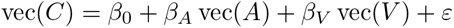

where *A*, *V*, and *C* denote the auditory-only, visual-only, and congruent audiovisual conditions, respectively, and vec(·) denotes vectorization of a 10 × 10 similarity matrix. Model estimation followed the same leave-one-run-out cross-validation procedure as our main RSA. On each fold, regression coefficients were estimated from the cells of the eight training folds and applied to the held-out fold to compute predicted values and residuals. Per-fold residual matrices were averaged across folds to yield a single cross-validated residual matrix per participant and ROI. Cells that could not be computed for a given fold due to trial exclusions remained missing and were dropped from both estimation and evaluation.

From this model we derived three quantities: (1) the regression coefficients *β_A_*and *β_V_*, indexing the contribution of each unimodal condition to the congruent audiovisual similarity; (2) the cross-validated *R*^2^ quantifying the proportion of variance in the congruent audiovisual matrix explained by the unimodal matrices on held-out folds; and (3) the residual congruent audiovisual MSPS, computed as the mean diagonal minus mean off-diagonal of the cross-validated residual matrix, indexing movie-specific pattern similarity in the congruent audiovisual condition that was not accounted for by the unimodal conditions.

### Whole-brain searchlight analyses

For exploratory analyses across the whole brain, we performed the cross-validated RSA approach in searchlights centered on every brain voxel. The searchlight analysis was implemented using BrainIAK (Brain Imaging Analysis Kit; [76]) and parallelized across voxels using the Message Passing Interface (MPI). For each participant, we swept a cubic searchlight (radius = 3 voxels) throughout the whole-brain mask. Searchlights were required to contain at least 80% of voxels within the participant’s whole-brain mask to be included in the analysis. At each searchlight location, we performed our RSA procedure for each condition (auditory-only, visual-only, congruent audiovisual, and incongruent audiovisual) as well as for crossmodal transfer. We also computed participant-level conjunctive contrast maps to assess multisensory facilitation (congruent audiovisual greater both unisensory conditions) and the disruptive effect of audiovisual incongruence (incongruent audiovisual less than mean of unisensory conditions). The resulting correlation coefficient maps were Fisher z-transformed and aligned into MNI152 standard space using ANTs [61] with linear interpolation for group-level analysis. Across participants, statistical maps were computed for each condition and contrast using FSL randomise with threshold-free cluster enhancement (TFCE; [74]).

### Statistical analyses

Statistical analyses were conducted in Python (version 3.7.9) using SciPy [77] (version 1.7.3), Statsmodels [78] (version 0.14.1), and custom code. All ROI tests used non-parametric sign-flipping randomization (10,000 iterations) with *α* = 0.05. Univariate GLM effects were evaluated using two-tailed tests; RSA-based pattern similarity was evaluated using one-tailed (greater than) tests given a priori hypotheses of movie-specific representational distinctiveness. Post-hoc pairwise comparisons between conditions used one-tailed randomization tests on the within-participant differences. FDR correction for multiple comparisons was applied separately across anatomical divisions of the ROIs: anterior vs. posterior hippocampus; left vs. right hippocampus, within each of the whole, anterior, and posterior hippocampus; hippocampal subfields (CA1, CA2/3, DG, and Subiculum); MTL cortex (PHC, PRC, and EC); and Harvard-Oxford cortical ROIs (OP, LOC, HG, and pSTG). Empirical null distributions and the corresponding observed group statistics for the multivariate results are shown in Figure S10. For whole-brain analyses (GLM and RSA searchlight), we used randomise in FSL (10,000 iterations; [74]) with TFCE correction to generate statistical maps.

### Data and code availability

The raw and preprocessed neuroimaging data will be released publicly upon acceptance. The task code is publicly available at https://github.com/omriraccah/multimem_fMRI_expt. The analysis code is publicly available at https://github.com/omriraccah/multimem_fMRI_analysis.

## Author contributions

O.R. and N.B.T-B. conceived of the study. O.R., Y.Z., and N.B.T-B. designed the experiment. O.R. and Y.Z. implemented the experiment and collected the data. O.R. and A.A. analyzed the data with support from Y.Z. O.R. and N.B.T-B. wrote the manuscript. All authors edited the article. N.B.T-B. provided supervision for all aspects of the study.

## Acknowledgments

O.R. was supported by a NIH Ruth L. Kirschstein Postdoctoral Individual National Research Service Award from the National Eye Institute (F32 EY035941). N.B.T-B. was supported by an NIH grant (R01 MH069456) and the Canadian Institute for Advanced Research. This study was supported by BrainWorks (RRID:SCR_024556) at the Center for Neurocognition and Behavior in the Wu Tsai Institute, Yale University

We thank Roeland Hancock, Alexander Forrence, Irene Zhou, Erica Busch, Lillian Behm, and Juliana Trach for their help with data collection. We also thank all Turk-Browne lab members for their extensive feedback and support.

## Supplementary Material

**Table S1:**
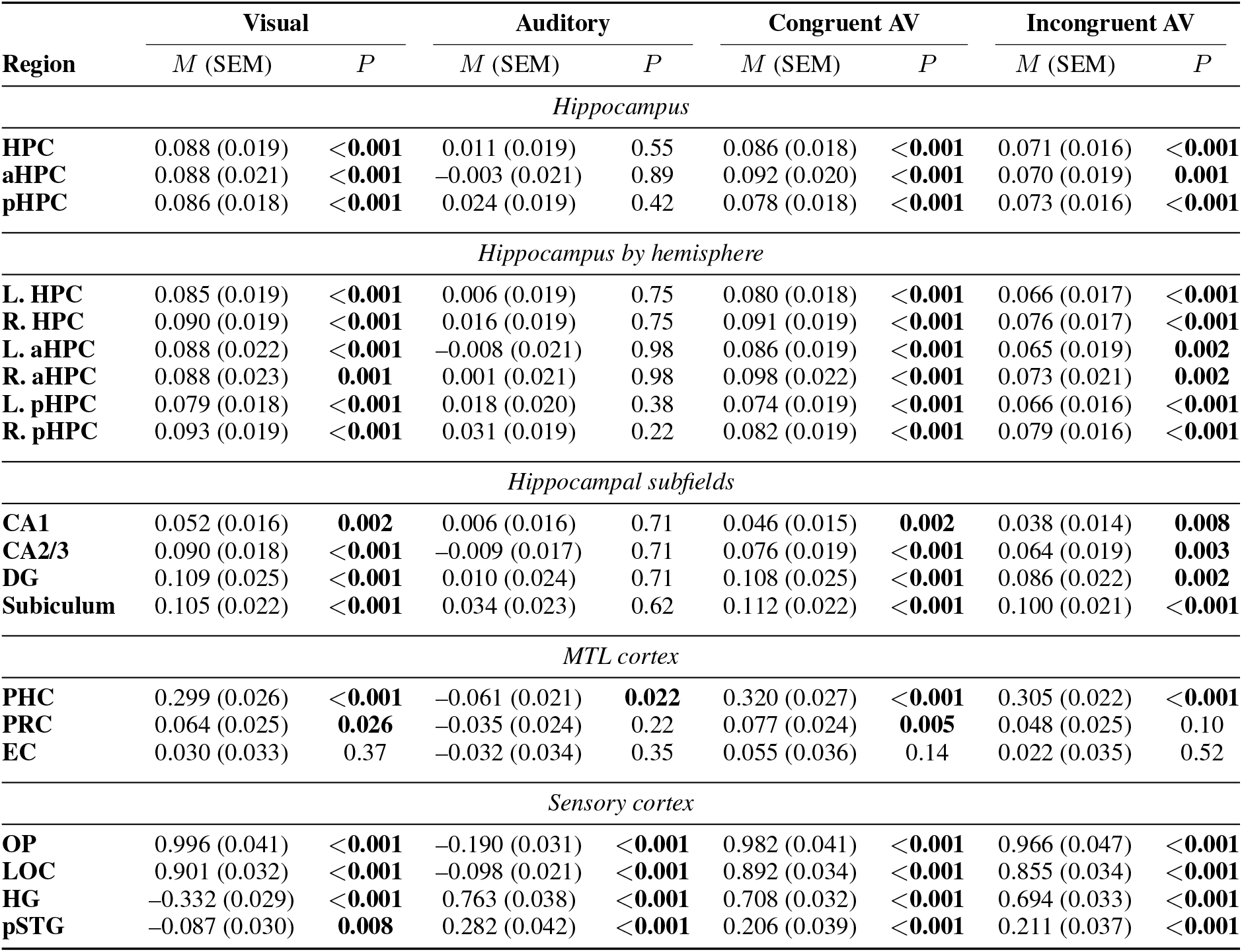
Univariate activation relative to baseline. *M* = mean *β* coefficient across participants. SEM = standard error of the mean. *P* = FDR-corrected statistical significance, bolded if *P <* 0.05.

**Table S2:** List of peak activations in the whole-brain GLM analysis. Significant clusters were identified using threshold-free cluster enhancement (TFCE) at *P <* 0.05. Bidirectional contrasts are reported in both directions (auditory vs. visual; congruent vs. incongruent). Anatomical labels were assigned to each peak using the Harvard-Oxford atlas [71]. For each cluster, the peak and any local maxima at least 8 mm apart (labeled a, b, c) are shown. Clusters smaller than 20 voxels are omitted.

| Cluster | Region | Hem. | MNI (*x, y, z*) | Peak *t* | Size (mm<sup>3</sup>) |
| --- | --- | --- | --- | --- | --- |
| *Auditory > visual* |  |  |  |  |  |
| 1 | Superior temporal gyrus (posterior) | R | 46, −19, −1 | 20.52 | 317530 |
| 1a | Heschl's gyrus | L | −49, −22, 9 | 20.04 |  |
| 1b | Heschl's gyrus | R | 47, −16, 3 | 19.11 |  |
| 1c | Planum temporale | L | −51, −31, 7 | 18.83 |  |
| 2 | Unlabeled | L | −30, −95, −30 | 5.61 | 3936 |
| 2a | Unlabeled | L | −35, −88, −36 | 5.53 |  |
| 2b | Unlabeled | L | −51, −77, −36 | 5.30 |  |
| 2c | Unlabeled | L | −41, −85, −31 | 5.08 |  |
| 3 | Postcentral gyrus | L | −38, −24, 40 | 5.60 | 44 |
| 4 | Precentral gyrus | R | 58, −3, 47 | 5.02 | 460 |
| *Visual > auditory* |  |  |  |  |  |
| 1 | Thalamus | L | −20, −31, 0 | 25.34 | 877945 |
| 1a | Occipital fusiform gyrus | L | −25, −84, −11 | 23.37 |  |
| 1b | Lingual gyrus | L | −7, −90, −8 | 22.07 |  |

| Cluster | Region | Hem. | MNI ($x, y, z$) | Peak $t$ | Size (mm<sup>3</sup>) |
| --- | --- | --- | --- | --- | --- |
| 1c | Thalamus | R | 22, -31, 0 | 22.01 |  |
| *Multisensory facilitation (congruent > auditory ∩ congruent > visual)* |  |  |  |  |  |
| 1 | Thalamus | R | 15, -26, -3 | 10.68 | 5947 |
| 1a | Thalamus | L | -18, -26, -5 | 10.27 |  |
| 1b | Thalamus | R | 14, -27, 4 | 7.85 |  |
| 1c | Brain-stem | L | -6, -34, -6 | 6.70 |  |
| 2 | Parietal operculum cortex | L | -45, -31, 18 | 6.62 | 5427 |
| 2a | Parietal operculum cortex | L | -51, -37, 28 | 6.44 |  |
| 2b | Parietal operculum cortex | L | -45, -34, 23 | 6.27 |  |
| 2c | Parietal operculum cortex | L | -49, -28, 18 | 5.98 |  |
| 3 | Parietal operculum cortex | R | 39, -27, 20 | 6.27 | 5706 |
| 3a | Parietal operculum cortex | R | 49, -28, 21 | 5.44 |  |
| 3b | Parietal operculum cortex | R | 61, -32, 22 | 4.87 |  |
| 3c | Planum temporale | R | 45, -38, 16 | 4.17 |  |
| 4 | Insular cortex | R | 37, -9, -5 | 5.32 | 922 |
| 4a | Insular cortex | R | 37, 2, -19 | 4.06 |  |
| 4b | Insular cortex | R | 38, 1, -10 | 3.60 |  |
| 4c | Temporal pole | R | 27, 4, -23 | 3.51 |  |
| 5 | Insular cortex | L | -35, -8, 13 | 4.85 | 1434 |
| 5a | Insular cortex | L | -36, -17, -7 | 4.55 |  |
| 5b | Insular cortex | L | -38, -8, -12 | 4.55 |  |
| 5c | Insular cortex | L | -37, -15, -7 | 4.30 |  |
| 6 | Supramarginal gyrus (posterior) | L | -53, -43, 13 | 3.63 | 376 |
| 6a | Planum temporale | L | -48, -46, 19 | 2.47 |  |
| 7 | Brain-stem | L | -11, -31, -24 | 3.59 | 54 |
| 8 | Temporal pole | L | -34, 2, -17 | 3.55 | 159 |
| 8a | Temporal pole | L | -32, 6, -28 | 2.91 |  |
| 9 | Frontal orbital cortex | R | 36, 31, -25 | 3.44 | 285 |
| 10 | Insular cortex | R | 30, -29, 8 | 3.37 | 70 |
| 11 | Frontal orbital cortex | R | 52, 29, -20 | 3.32 | 77 |
| 12 | Precentral gyrus | R | 61, -5, 24 | 3.06 | 306 |
| 12a | Postcentral gyrus | R | 62, -5, 15 | 2.39 |  |
| 13 | Angular gyrus | R | 61, -48, 14 | 2.55 | 182 |
| *Congruent > incongruent* |  |  |  |  |  |
| 1 | Paracingulate gyrus | L | -6, 50, -2 | 7.01 | 3382 |
| 1a | Paracingulate gyrus | L | -7, 42, -10 | 5.90 |  |
| 1b | Frontal medial cortex | L | -9, 51, -6 | 5.84 |  |
| 1c | Frontal medial cortex | R | 9, 40, -15 | 5.41 |  |
| 2 | Lateral occipital cortex (inferior) | R | 57, -68, -6 | 4.54 | 38 |
| *Incongruent > congruent* |  |  |  |  |  |
| 1 | Superior parietal lobule | L | -38, -57, 45 | 6.28 | 1381 |
| 2 | Inferior frontal gyrus (pars opercularis) | L | -41, 15, 24 | 6.07 | 4278 |
| 2a | Middle frontal gyrus | L | -53, 18, 29 | 5.60 |  |
| 2b | Inferior frontal gyrus (pars triangularis) | L | -54, 24, 23 | 5.37 |  |
| 2c | Middle frontal gyrus | L | -36, 5, 33 | 5.19 |  |
| 3 | Middle frontal gyrus | R | 52, 22, 29 | 5.78 | 512 |
| 3a | Middle frontal gyrus | R | 42, 23, 22 | 5.60 |  |

**Figure S1:**
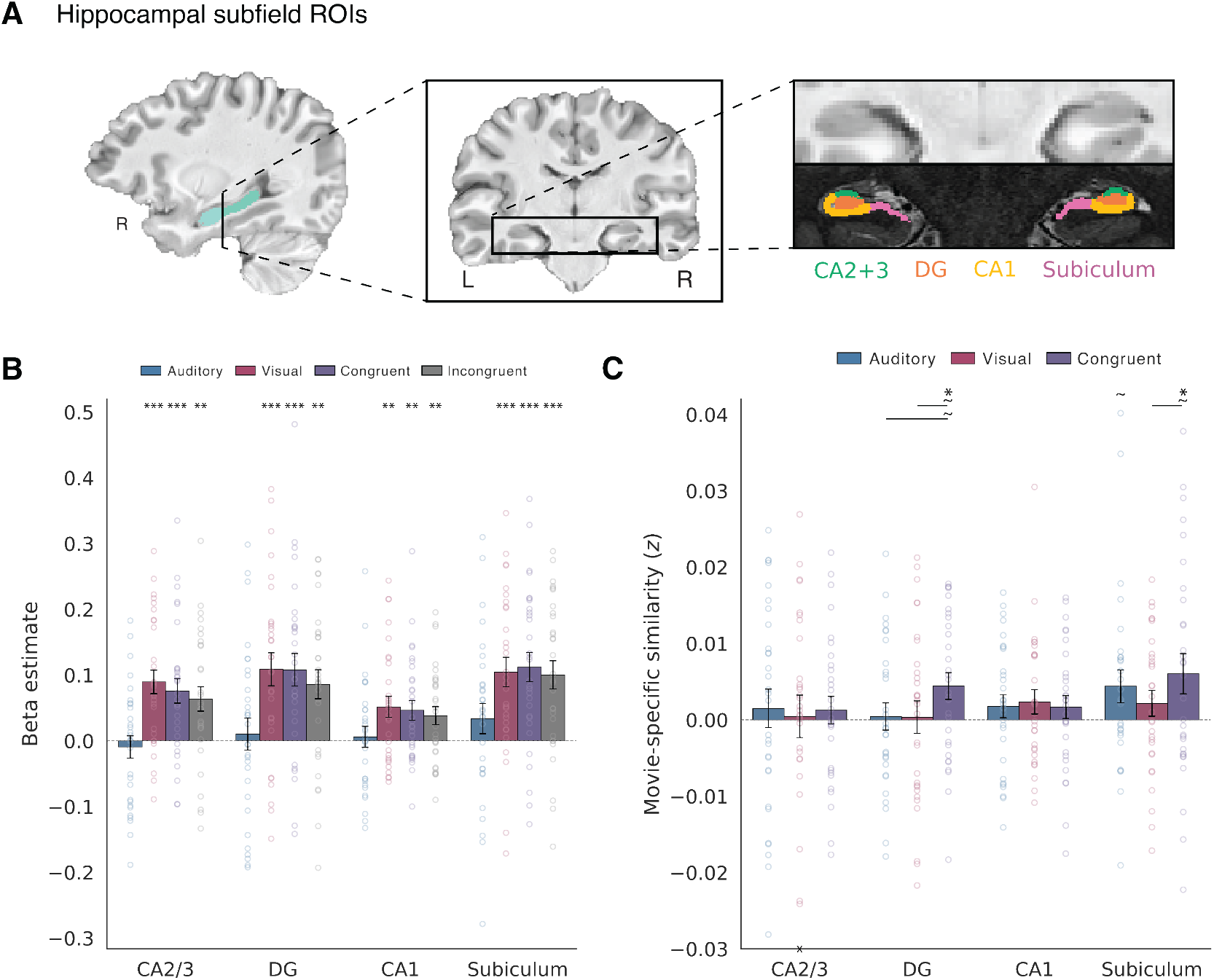
Univariate and multivariate results in human hippocampal subfields. **(A)** Hippocampal subfields (CA2/3, DG, CA1, and subiculum) were segmented using the ASHS toolbox [68] trained on an atlas of manual 3T segmentations [70, 69]. **(B)** Univariate GLM results in hippocampal subfields. **(C)** Multivariate RSA results in hippocampal subfields. Error bars indicate SEM. ∼ *P <* 0.1, ^∗^*P <* 0.05, ^∗∗^*P <* 0.01, ^∗∗∗^*P <* 0.001.

**Table S3:** Multivariate pattern similarity relative to baseline. *M* = mean movie-specific pattern similarity across participants. SEM = standard error of the mean. *P* = FDR-corrected statistical significance, bolded if *P <* 0.05.

| Region | Visual |  | Auditory |  | Congruent AV |  |
| --- | --- | --- | --- | --- | --- | --- |
|  | *M* (SEM) | *P* | *M* (SEM) | *P* | *M* (SEM) | *P* |
| *Hippocampus* |  |  |  |  |  |  |
| **HPC** | 0.002 (0.001) | 0.11 | 0.002 (0.001) | **0.017** | 0.004 (0.001) | **0.001** |
| **aHPC** | −0.000 (0.002) | 0.51 | 0.004 (0.001) | **0.007** | 0.003 (0.001) | **0.012** |
| **pHPC** | 0.003 (0.001) | **0.031** | 0.000 (0.001) | 0.46 | 0.005 (0.002) | **0.003** |
| *Hippocampus by hemisphere* |  |  |  |  |  |  |
| **L. HPC** | 0.001 (0.002) | 0.36 | 0.002 (0.001) | 0.072 | 0.004 (0.002) | **0.026** |
| **R. HPC** | 0.002 (0.002) | 0.24 | 0.003 (0.001) | 0.058 | 0.005 (0.001) | **0.001** |
| **L. aHPC** | −0.000 (0.002) | 0.54 | 0.003 (0.002) | 0.066 | 0.002 (0.002) | 0.20 |
| **R. aHPC** | −0.000 (0.002) | 0.54 | 0.004 (0.002) | **0.013** | 0.005 (0.002) | **0.010** |
| **L. pHPC** | 0.001 (0.002) | 0.33 | 0.000 (0.002) | 0.55 | 0.006 (0.002) | **0.011** |
| **R. pHPC** | 0.004 (0.002) | **0.039** | −0.000 (0.002) | 0.55 | 0.004 (0.002) | **0.036** |
| *Hippocampal subfields* |  |  |  |  |  |  |
| **CA1** | 0.002 (0.002) | 0.19 | 0.002 (0.002) | 0.24 | 0.002 (0.002) | 0.19 |
| **CA2/3** | 0.000 (0.003) | 0.43 | 0.002 (0.003) | 0.36 | 0.001 (0.002) | 0.24 |
| **DG** | 0.000 (0.002) | 0.43 | 0.000 (0.002) | 0.40 | 0.004 (0.002) | **0.031** |
| **Subiculum** | 0.002 (0.002) | 0.19 | 0.004 (0.002) | 0.088 | 0.006 (0.003) | **0.031** |
| *MTL cortex* |  |  |  |  |  |  |
| **PHC** | 0.045 (0.003) | <**0.001** | 0.006 (0.002) | **0.002** | 0.045 (0.003) | <**0.001** |
| **PRC** | 0.001 (0.002) | 0.33 | 0.001 (0.002) | 0.54 | 0.003 (0.002) | **0.027** |
| **EC** | −0.000 (0.003) | 0.55 | −0.002 (0.003) | 0.69 | 0.002 (0.003) | 0.32 |
| *Sensory cortex* |  |  |  |  |  |  |
| **OP** | 0.061 (0.008) | <**0.001** | 0.006 (0.004) | 0.084 | 0.071 (0.006) | <**0.001** |
| **LOC** | 0.090 (0.020) | <**0.001** | 0.021 (0.008) | **0.004** | 0.085 (0.009) | <**0.001** |
| **HG** | 0.009 (0.002) | <**0.001** | 0.083 (0.005) | <**0.001** | 0.088 (0.005) | <**0.001** |
| **pSTG** | 0.025 (0.005) | <**0.001** | 0.053 (0.004) | <**0.001** | 0.067 (0.006) | <**0.001** |

**Table S4:** Multisensory effects. Multivariate evidence for multisensory facilitation (congruent *>* visual; congruent *>* auditory) and crossmodal transfer. (Δ)*M* = (difference in) mean movie-specific pattern similarity across participants. SEM = standard error of the mean. *P* = FDR-corrected statistical significance, bolded if *<* 0.05. Facilitation was tested only in regions where congruent numerically exceeded both unisensory conditions (— = not tested).

| Region | Congruent > Visual |  | Congruent > Auditory |  | Crossmodal Transfer |  |
| --- | --- | --- | --- | --- | --- | --- |
| | $\Delta M$ (SEM) | $P$ | $\Delta M$ (SEM) | $P$ | $M$ (SEM) | $P$ |
| *Hippocampus* |  |  |  |  |  |  |
| **HPC** | 0.003 (0.001) | **0.029** | 0.002 (0.002) | 0.15 | 0.002 (0.001) | **0.003** |
| **aHPC** | — | — | — | — | 0.003 (0.001) | **0.003** |
| **pHPC** | 0.002 (0.001) | 0.084 | 0.005 (0.002) | **0.013** | 0.001 (0.001) | 0.22 |
| *Hippocampus by hemisphere* |  |  |  |  |  |  |
| **L. HPC** | 0.003 (0.002) | 0.099 | 0.002 (0.003) | 0.23 | 0.001 (0.001) | 0.26 |
| **R. HPC** | 0.003 (0.001) | 0.074 | 0.002 (0.002) | 0.23 | 0.003 (0.001) | **0.006** |
| **L. aHPC** | — | — | — | — | 0.001 (0.001) | 0.20 |
| **R. aHPC** | — | — | — | — | 0.004 (0.002) | **0.016** |
| **L. pHPC** | 0.005 (0.002) | **0.009** | 0.005 (0.003) | **0.040** | 0.000 (0.001) | 0.49 |
| **R. pHPC** | — | — | — | — | 0.002 (0.001) | 0.16 |
| *Hippocampal subfields* |  |  |  |  |  |  |
| **CA1** | — | — | — | — | 0.000 (0.001) | 0.42 |
| **CA2/3** | — | — | — | — | 0.002 (0.001) | 0.14 |
| **DG** | 0.004 (0.003) | 0.065 | 0.004 (0.002) | 0.074 | 0.002 (0.001) | 0.091 |
| **Subiculum** | 0.004 (0.002) | 0.065 | 0.002 (0.004) | 0.34 | 0.002 (0.001) | 0.091 |
| *MTL cortex* |  |  |  |  |  |  |
| **PHC** | — | — | — | — | 0.008 (0.001) | **<0.001** |
| **PRC** | 0.002 (0.002) | 0.28 | 0.003 (0.003) | 0.25 | 0.002 (0.001) | 0.14 |
| **EC** | 0.002 (0.005) | 0.35 | 0.003 (0.004) | 0.25 | 0.001 (0.002) | 0.39 |
| *Sensory cortex* |  |  |  |  |  |  |
| **OP** | 0.010 (0.008) | 0.13 | 0.065 (0.006) | **<0.001** | 0.005 (0.002) | **0.020** |
| **LOC** | — | — | — | — | 0.017 (0.013) | **0.047** |
| **HG** | 0.079 (0.005) | **<0.001** | 0.005 (0.002) | **0.025** | 0.004 (0.001) | **0.004** |
| **pSTG** | 0.042 (0.005) | **<0.001** | 0.014 (0.005) | **0.006** | 0.009 (0.003) | **0.004** |

**Table S5:** Unisensory contributions to congruent audiovisual similarity. Cross-validated multiple regression modeling the congruent audiovisual similarity matrix as a function of the auditory-only and visual-only similarity matrices. *β_A_*, *β_V_* = auditory-only and visual-only regression coefficients. *R*^2^ = proportion of variance in the congruent audiovisual matrix explained by the unisensory matrices on held-out folds. Residual MSPS = movie-specific pattern similarity in the congruent audiovisual condition remaining after regressing out the unisensory contributions. *M* = mean across participants. SEM = standard error of the mean. *P* = FDR-corrected statistical significance, bolded if *<* 0.05. *R*^2^ was tested one-tailed against zero; *P* values approaching 1 therefore indicate cross-validated variance explained reliably below zero.

| Region | $\beta_A$ | | $\beta_V$ | | $R^2$ | | Residual MSPS | |
| --- | --- | --- | --- | --- | --- | --- | --- | --- |
| | $M$ (SEM) | $P$ | $M$ (SEM) | $P$ | $M$ (SEM) | $P$ | $M$ (SEM) | $P$ |
| *Hippocampus* |  |  |  |  |  |  |  |  |
| HPC | -0.008 (0.013) | 0.72 | 0.053 (0.021) | **0.003** | -0.005 (0.007) | 0.68 | 0.004 (0.001) | **0.002** |
| aHPC | 0.013 (0.014) | 0.36 | 0.016 (0.011) | 0.082 | -0.008 (0.002) | 1.00 | 0.003 (0.001) | **0.019** |
| pHPC | -0.019 (0.014) | 0.90 | 0.051 (0.021) | **0.005** | -0.008 (0.007) | 1.00 | 0.005 (0.002) | **0.005** |
| *Sensory cortex* |  |  |  |  |  |  |  |  |
| OP | 0.049 (0.019) | **0.009** | 0.344 (0.032) | **<0.001** | 0.108 (0.023) | **<0.001** | 0.044 (0.005) | **<0.001** |
| LOC | 0.039 (0.027) | 0.079 | 0.316 (0.041) | **<0.001** | 0.122 (0.028) | **<0.001** | 0.050 (0.006) | **<0.001** |
| HG | 0.565 (0.031) | **<0.001** | 0.075 (0.026) | **0.003** | 0.358 (0.040) | **<0.001** | 0.037 (0.002) | **<0.001** |
| pSTG | 0.283 (0.034) | **<0.001** | 0.113 (0.034) | **<0.001** | 0.108 (0.024) | **<0.001** | 0.045 (0.005) | **<0.001** |

**Figure S2:**
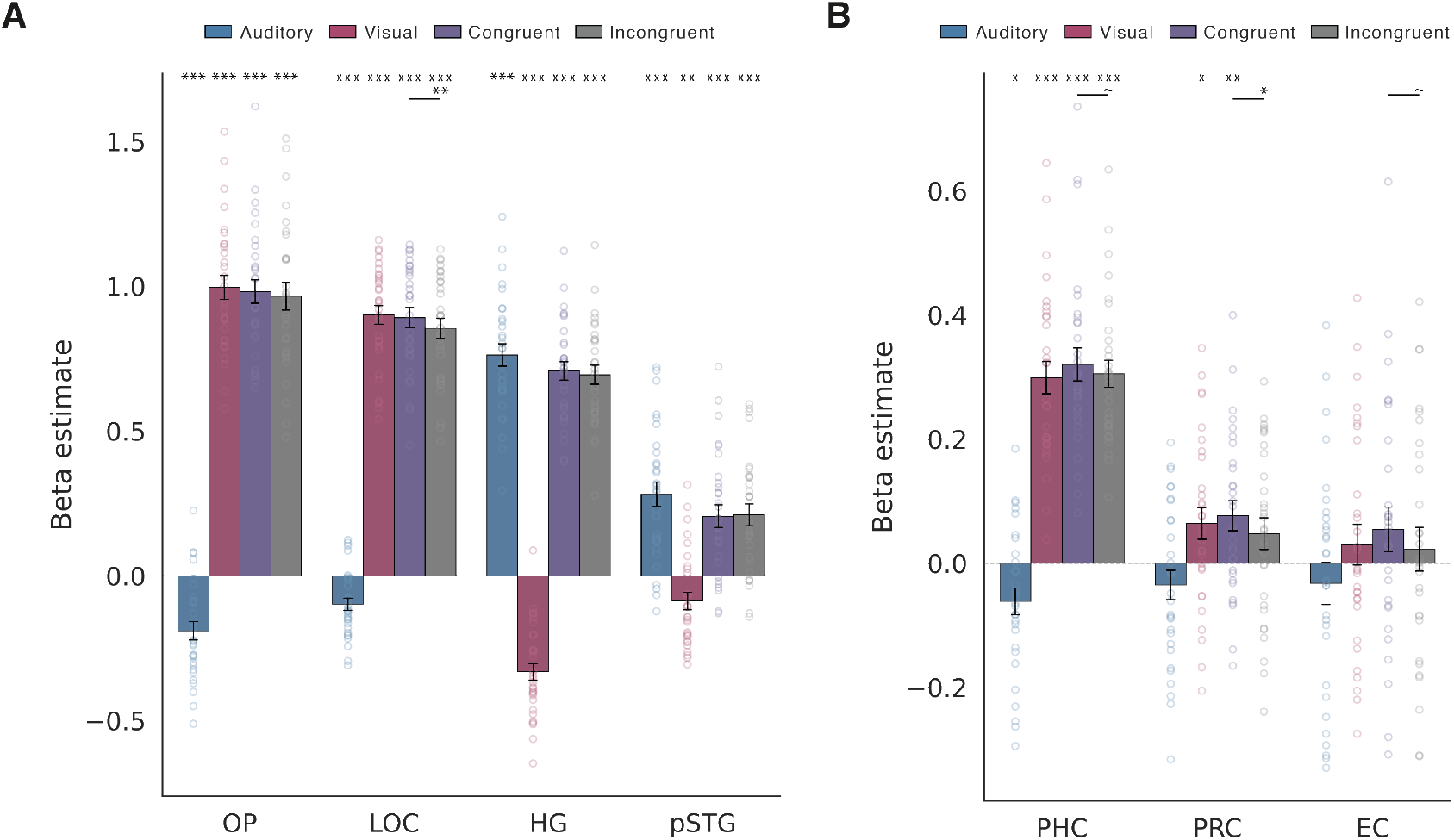
Univariate GLM results in sensory regions and MTL cortex. **(A)** Mean beta estimates for each condition in sensory cortical ROIs (OP, LOC, HG, pSTG). **(B)** Mean beta estimates for each condition in MTL cortical ROIs (PHC, PRC, EC). Individual participant data shown as circles; error bars indicate SEM. ∼ *P <* 0.1, ^∗^*P <* 0.05, ^∗∗^*P <* 0.01, ^∗∗∗^*P <* 0.001.

**Figure S3:**
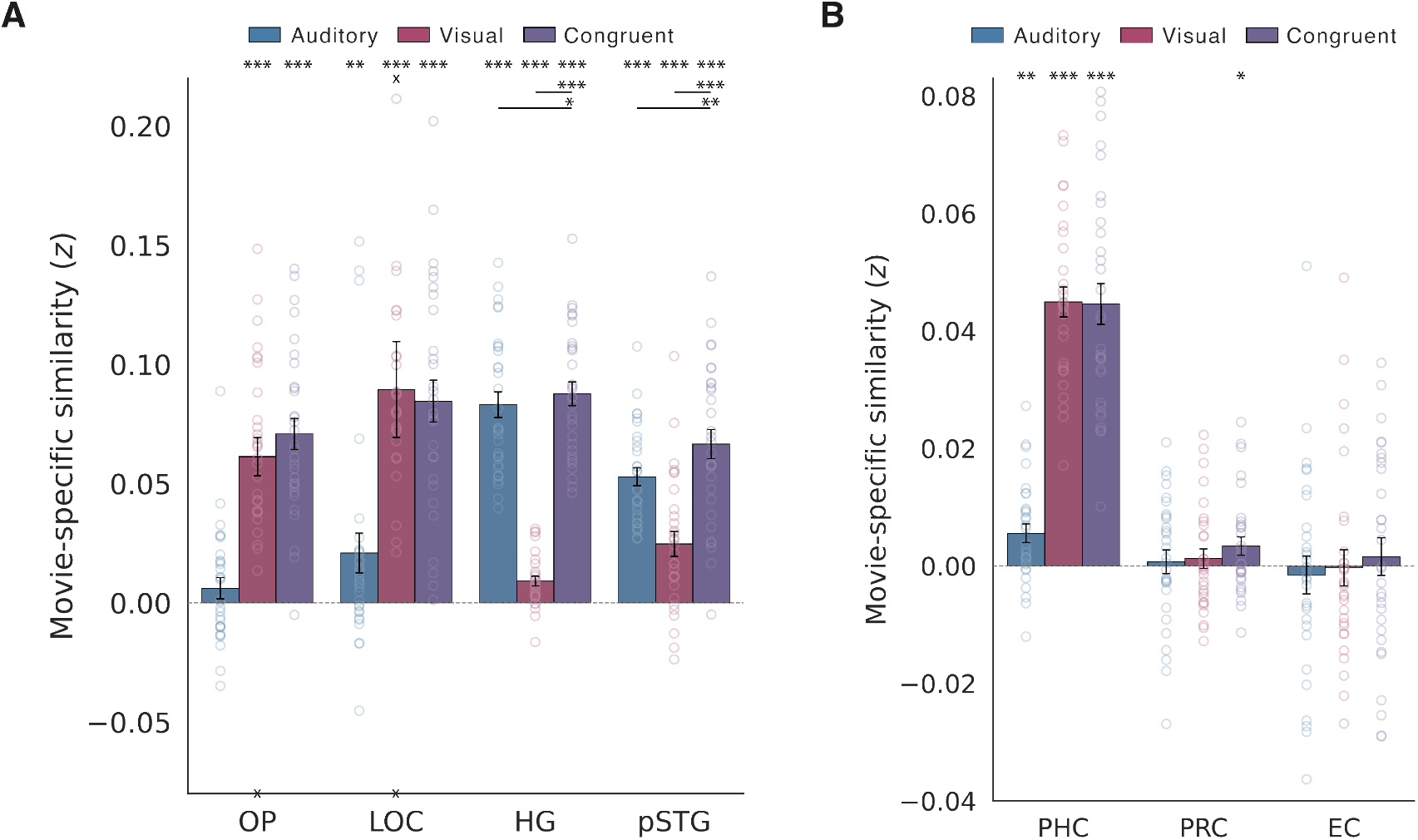
Multivariate RSA results in sensory regions and MTL cortex. **(A)** Movie-specific pattern similarity (MSPS) for each condition in sensory cortical ROIs (OP, LOC, HG, pSTG). HG and pSTG showed significant multisensory facilitation. **(B)** MSPS for each condition in MTL cortical ROIs (PHC, PRC, EC). Individual participant data shown as circles; error bars indicate SEM. × markers indicate individual data points outside the plotted y-axis range. ^∗^*P <* 0.05, ^∗∗^*P <* 0.01, ^∗∗∗^*P <* 0.001.

**Figure S4:**
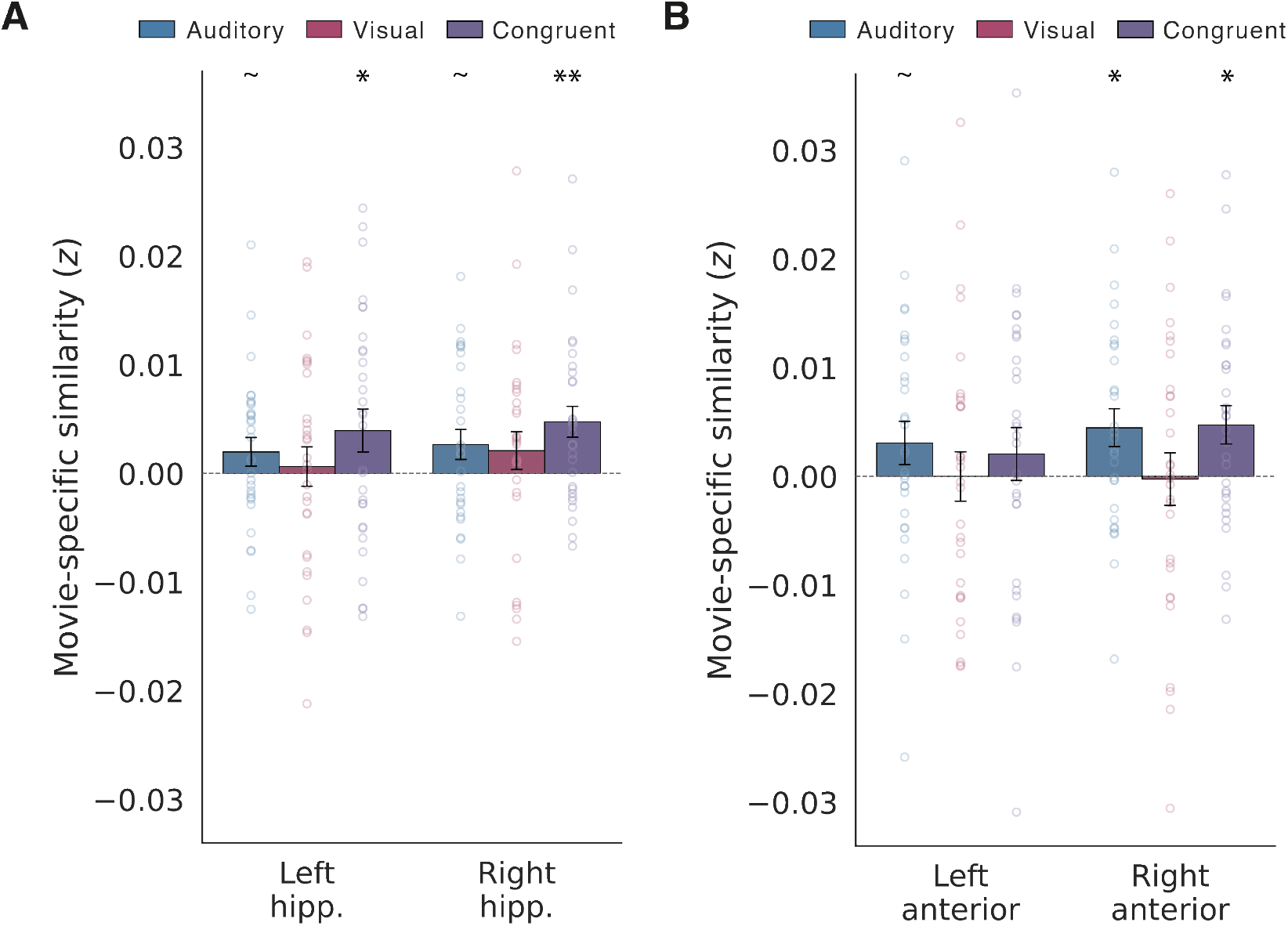
Condition-wise lateralization in the whole and anterior hippocampus. **(A)** MSPS for each condition in the left and right whole hippocampus. **(B)** MSPS for each condition in the left and right anterior hippocampus. Individual participant data shown as circles; error bars indicate SEM. ∼ *P <* 0.1, ^∗^*P <* 0.05, ^∗∗^*P <* 0.01.

**Figure S5:**
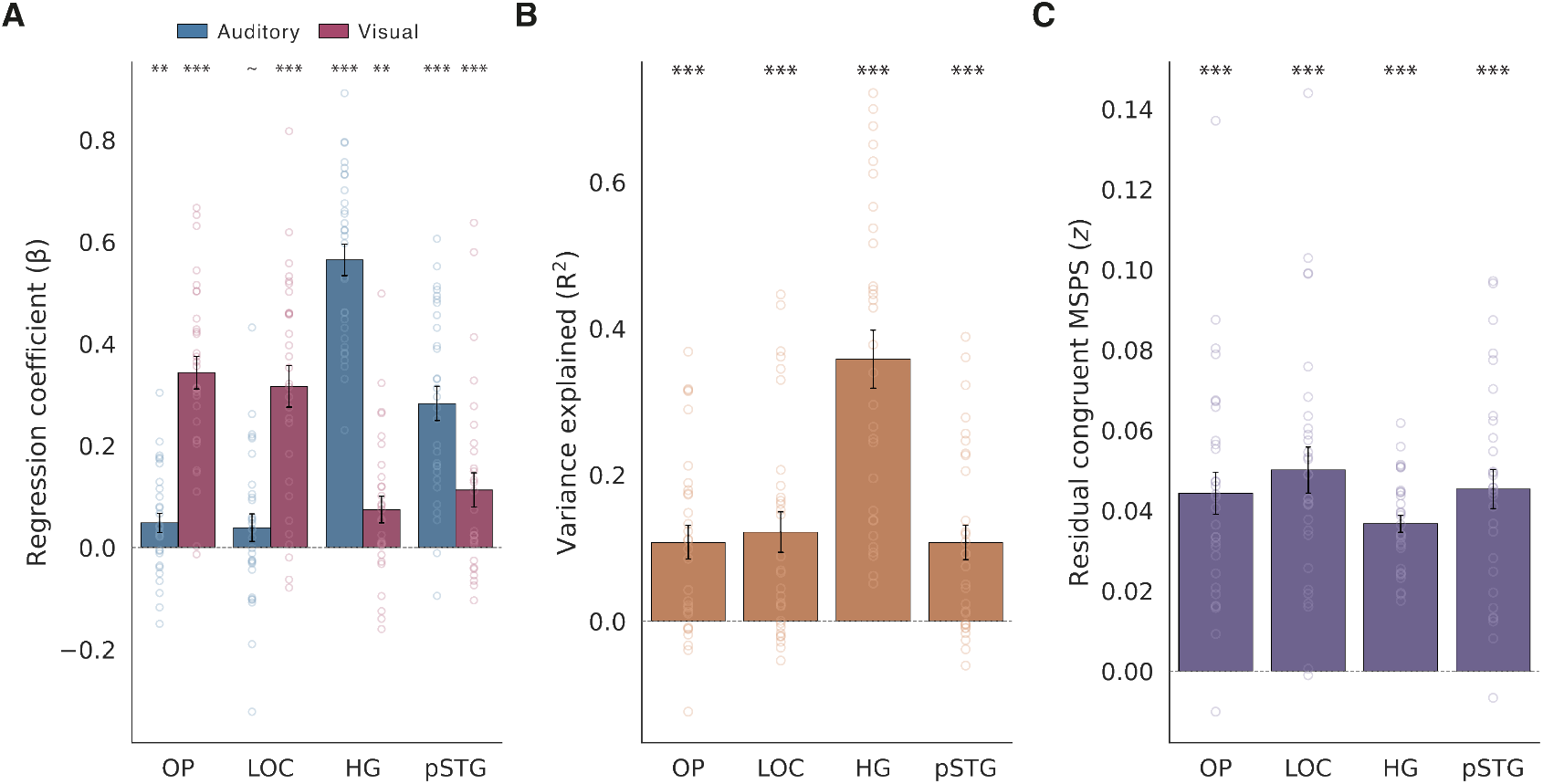
Conjunctive audiovisual coding in sensory regions. **(A)** Auditory and visual regression coefficients from the cross-validated multiple regression in sensory cortical ROIs (OP, LOC, HG, pSTG). **(B)** Variance explained (*R*^2^) by the unisensory matrices. **(C)** Residual congruent audiovisual MSPS after regressing out the auditory-only and visual-only contributions. Individual participant data shown as circles; error bars indicate SEM. × markers indicate individual data points outside the plotted y-axis range. ∼ *P <* 0.1, ^∗^*P <* 0.05, ^∗∗^*P <* 0.01, ^∗∗∗^*P <* 0.001.

**Figure S6:**
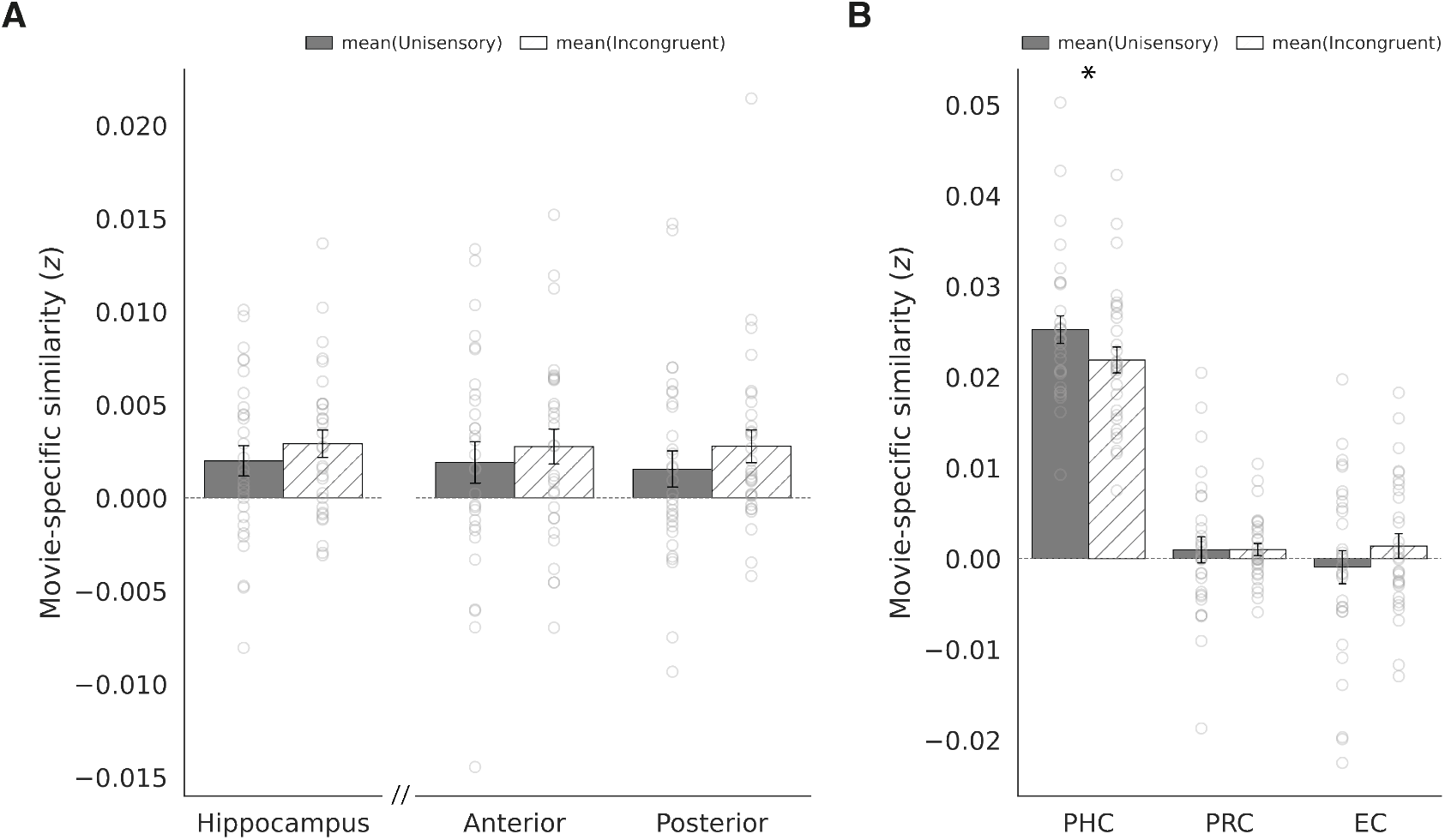
Unisensory versus incongruent MSPS in the hippocampus and MTL cortex. **(A)** Incongruent and unisensory (average of auditory-only and visual-only) MSPS in the whole hippocampus and anterior/posterior hippocampus. No significant differences were observed across these ROIs. **(B)** MSPS in MTL cortical ROIs (PHC, PRC, EC). PHC showed greater MSPS for unisensory relative to incongruent conditions, suggesting a catastrophic interaction between non-matching auditory and visual content. There were no significant differences in PRC or EC. Individual participant data shown as circles; error bars indicate SEM. ^∗^*P <* 0.05.

**Figure S7:**
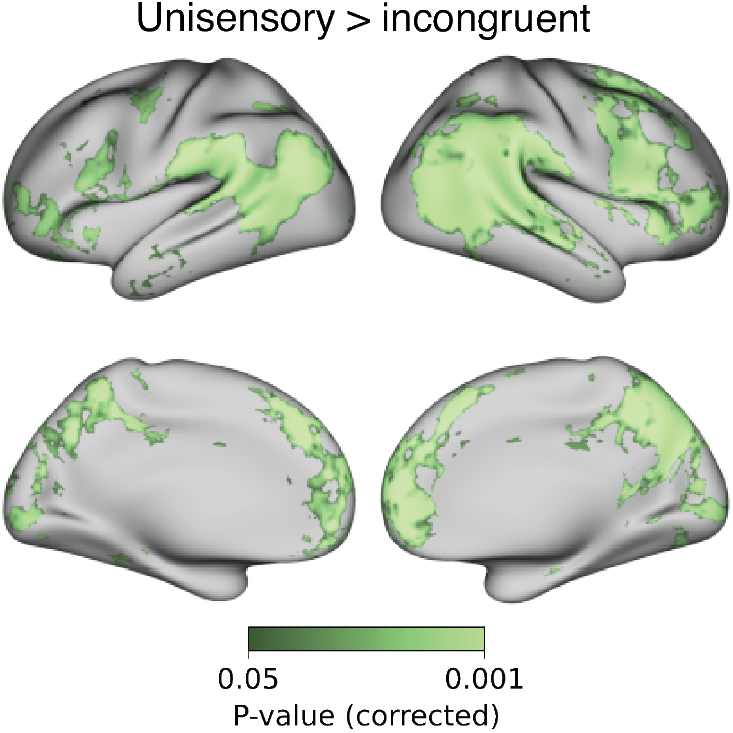
Whole-brain RSA searchlight of unisensory versus incongruent pattern similarity. MSPS was significantly reduced for incongruent relative to unisensory (average of auditory and visual) throughout widespread cortical sites, including the posterior STG, lateral occipital cortex, angular gyrus, posteromedial cortex, and the lateral and medial prefrontal cortex. A full list of peak coordinates is shown in Table S6. Corrected using TFCE at *P <* 0.05.

**Figure S8:**
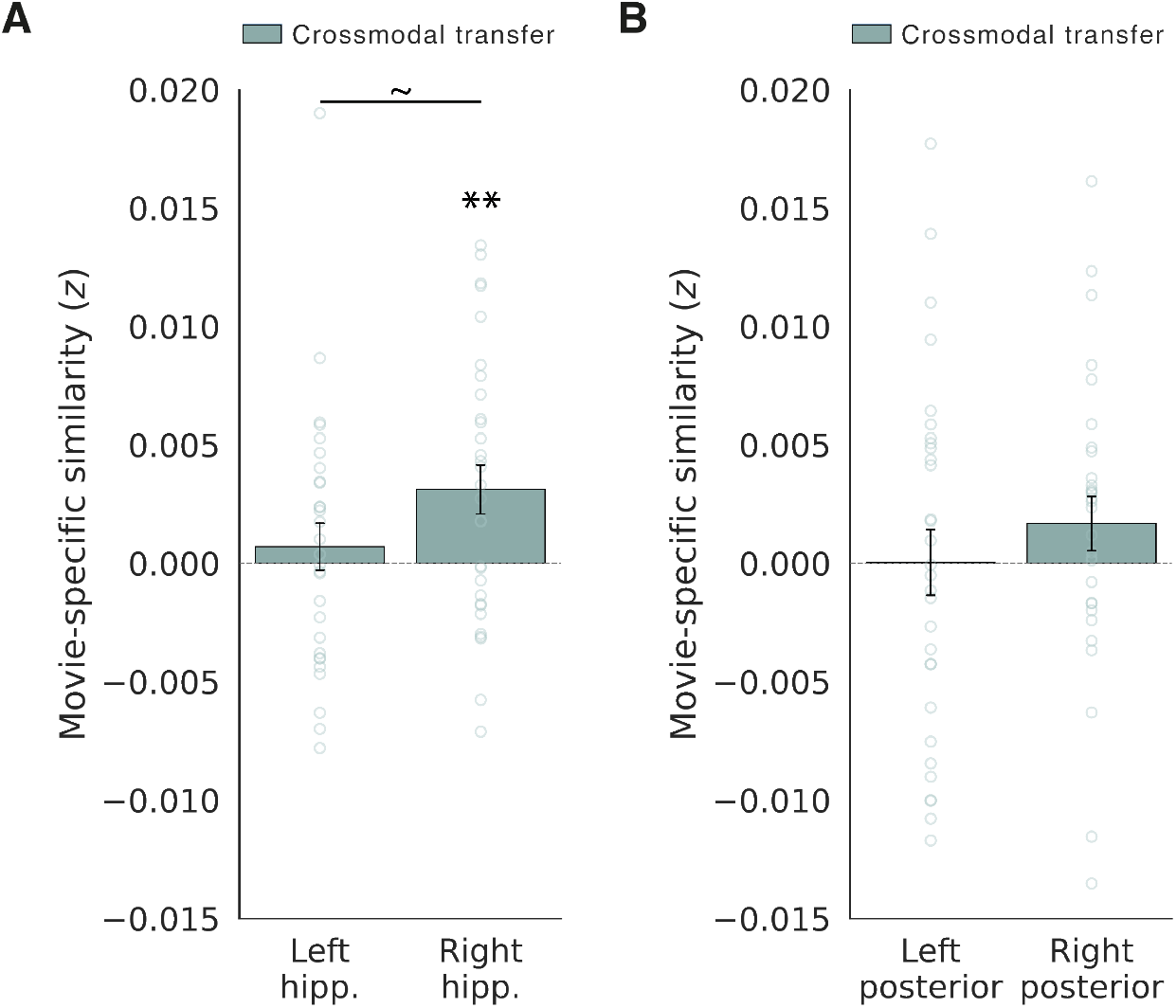
Crossmodal transfer lateralization in the whole and posterior hippocampus. **(A)** Crossmodal transfer MSPS in the left and right whole hippocampus. **(B)** Crossmodal transfer MSPS in the left and right posterior hippocampus. Individual participant data shown as circles; error bars indicate SEM. ∼ *P <* 0.1, ^∗^*P <* 0.05, ^∗∗^*P <* 0.01, ^∗∗∗^*P <* 0.001.

**Figure S9:**
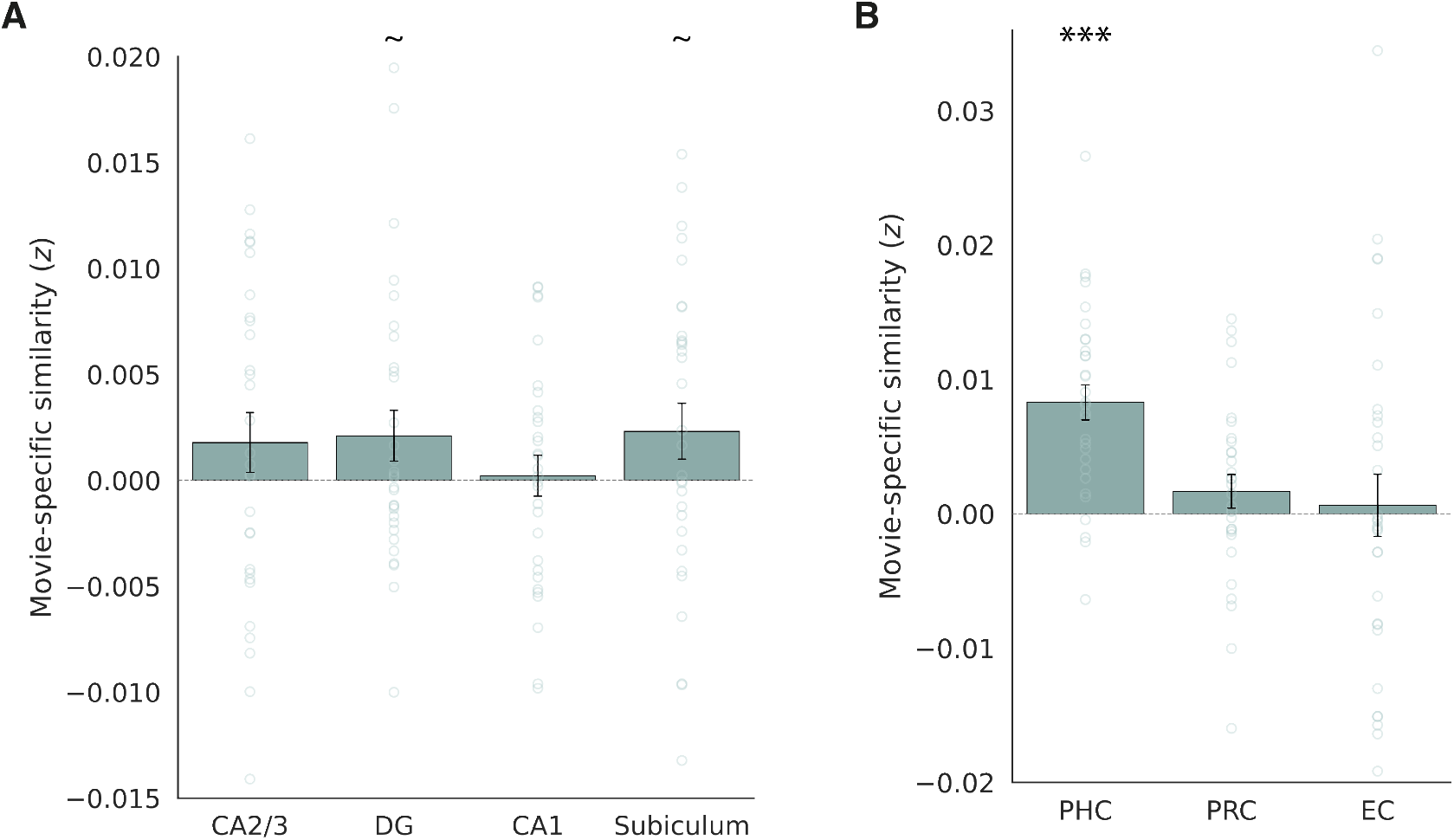
Crossmodal transfer in hippocampal subfields and MTL cortex. **(A)** Crossmodal transfer MSPS across hippocampal subfields (CA2/3, DG, CA1, subiculum). **(B)** Crossmodal transfer MSPS in MTL cortical ROIs (PHC, PRC, EC). Individual participant data shown as circles; error bars indicate SEM. ∼ *P <* 0.1, ^∗^*P <* 0.05, ^∗∗^*P <* 0.01, ^∗∗∗^*P <* 0.001.

**Figure S10:**
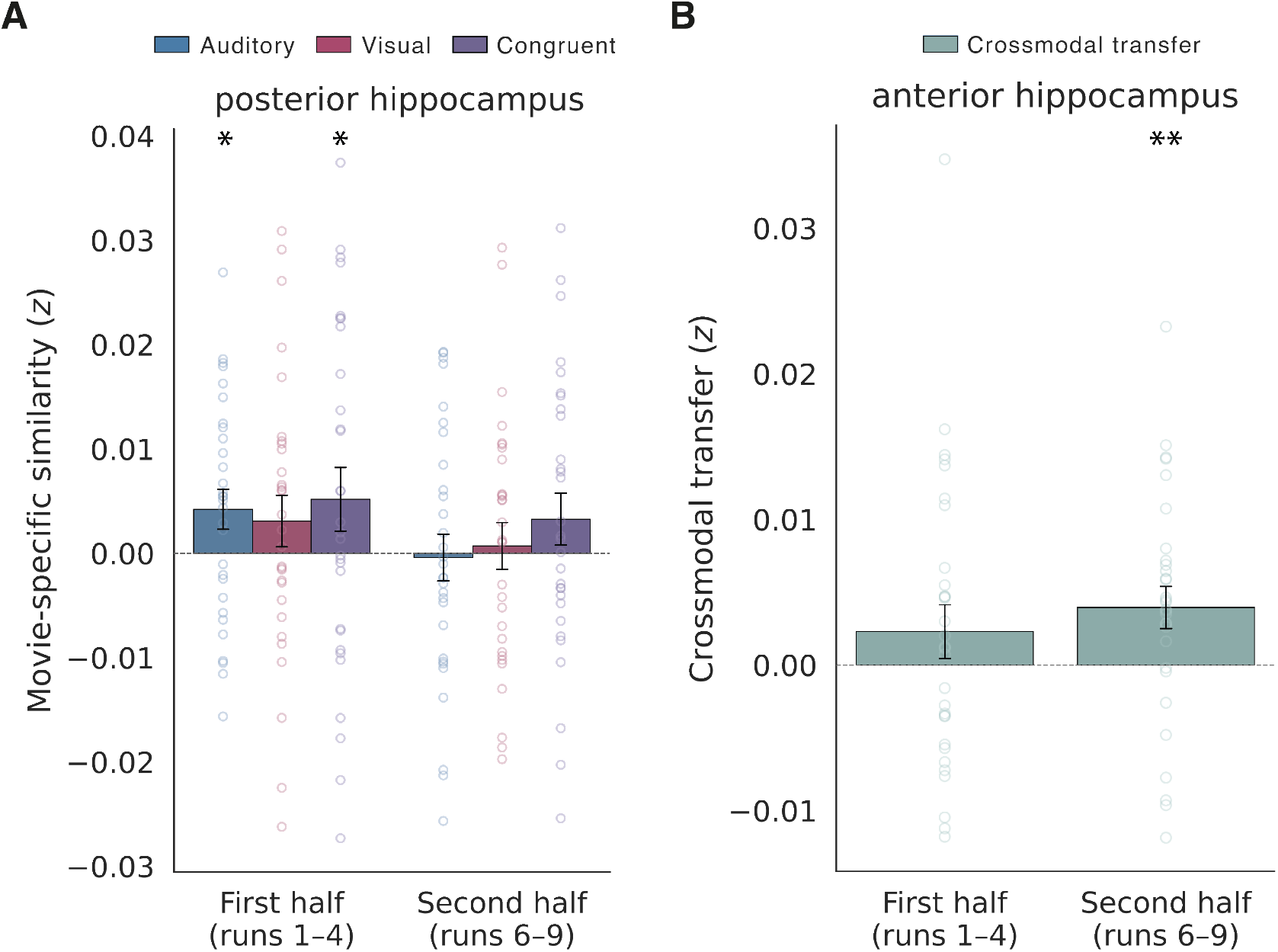
Split-half analysis of multisensory facilitation and crossmodal transfer. **(A)** MSPS for each condition in the posterior hippocampus, estimated separately within the first half (runs 1–4) and second half (runs 6–9) of the experiment. **(B)** Crossmodal transfer MSPS in the anterior hippocampus across the same two halves. Individual participant data shown as circles; error bars indicate SEM. ^∗^*P <* 0.05, ^∗∗^*P <* 0.01.

**Figure S11:**
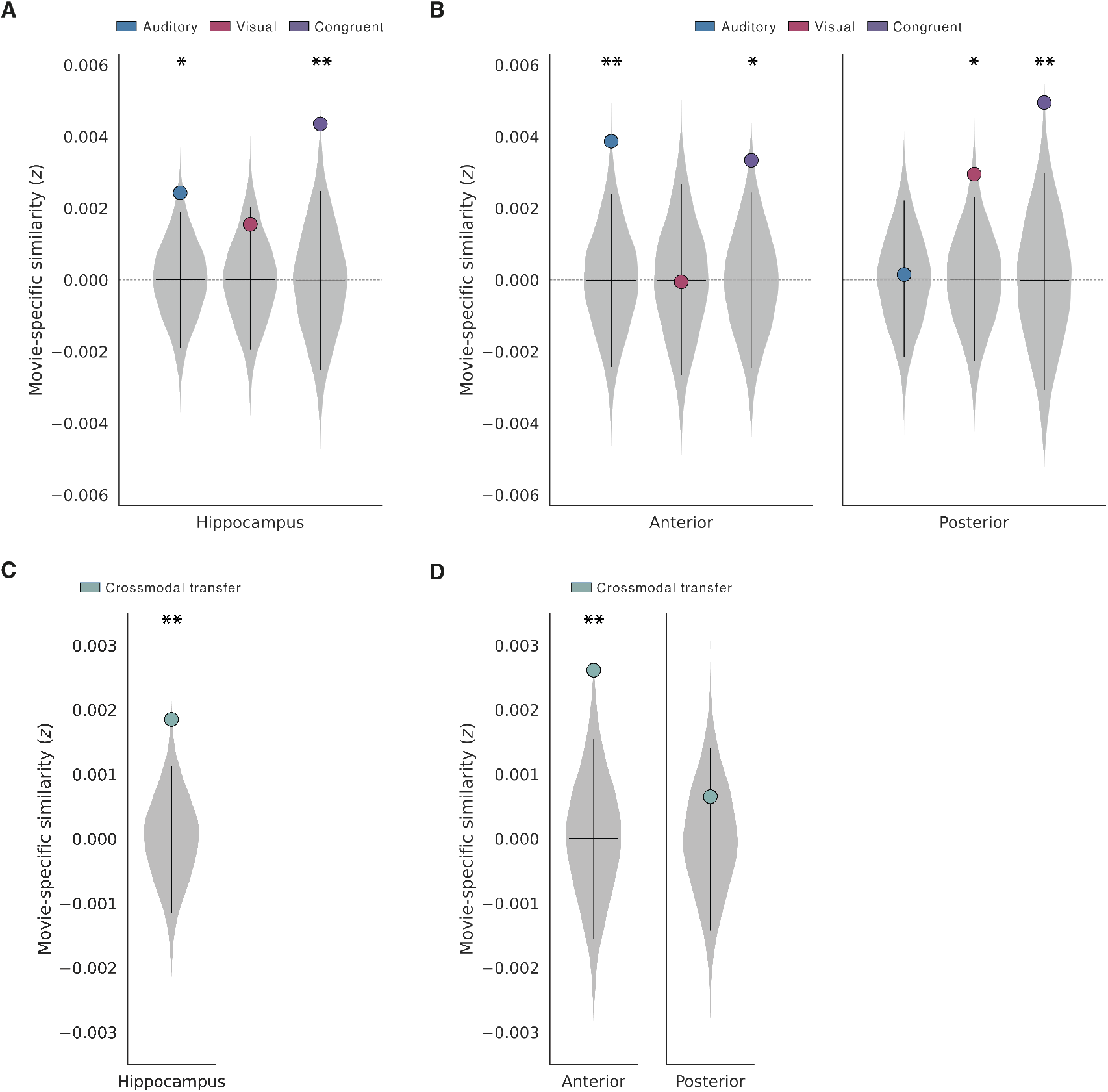
Permutation null distributions for multivariate effects. Mean MSPS across participants (colored circles) plotted against the empirical null distributions generated by the same sign-flipping randomization procedure used for statistical testing (10,000 iterations; grey violins). **(A)** MSPS for the auditory-only, visual-only, and congruent audiovisual conditions in the whole hippocampus. **(B)** The same, shown separately for the anterior and posterior hippocampus. **(C)** Crossmodal transfer MSPS in the whole hippocampus. **(D)** Crossmodal transfer MSPS in the anterior and posterior hippocampus. Grey violins show the kernel density of the null distribution for each ROI and condition. The vertical black bar spans the 5th to 95th percentile of the null and the horizontal black tick marks the null mean. Asterisks denote FDR-corrected significance. ^∗^*P <* 0.05, ^∗∗^*P <* 0.01, ^∗∗∗^*P <* 0.001.

**Table S6:**
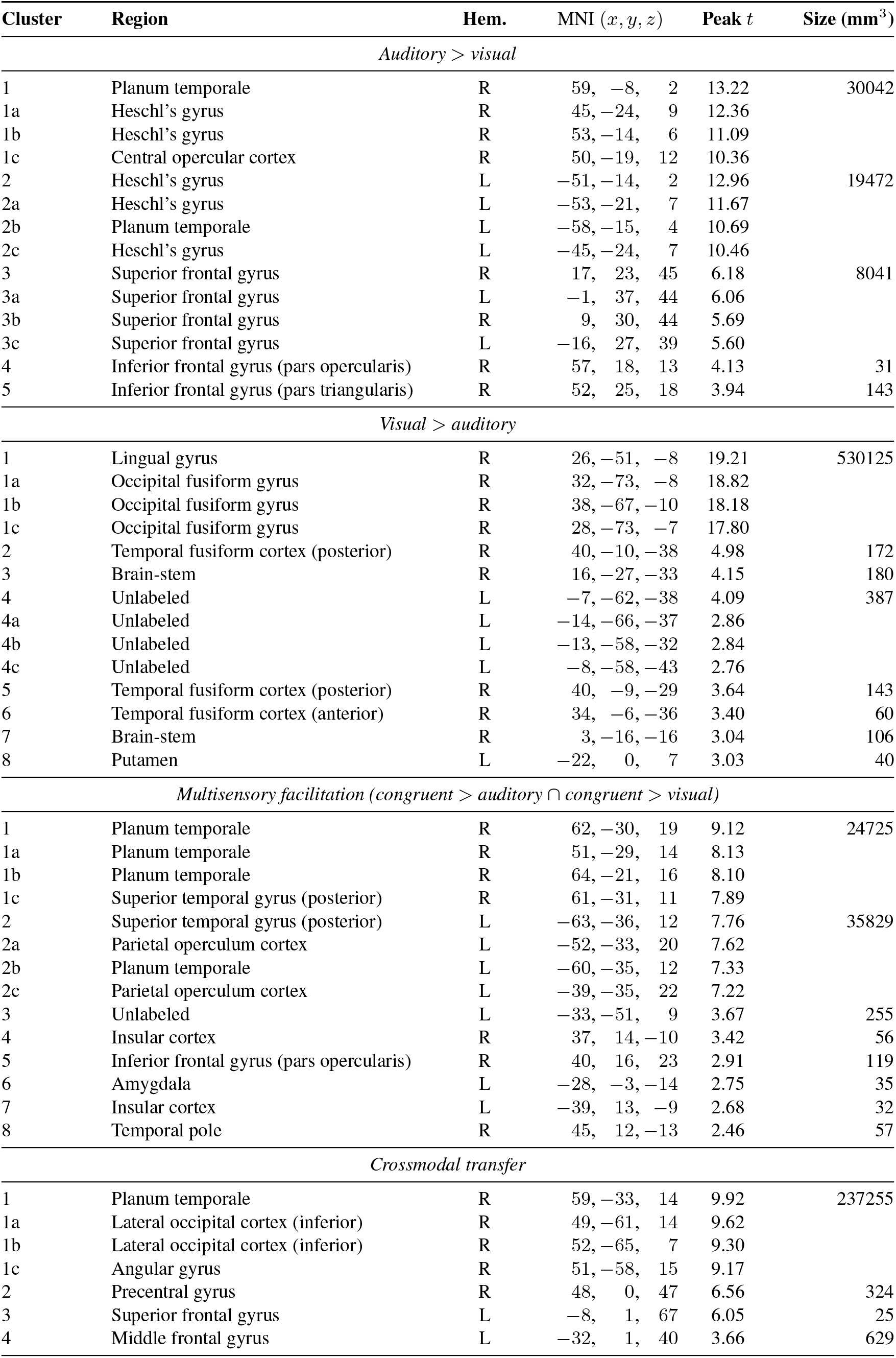
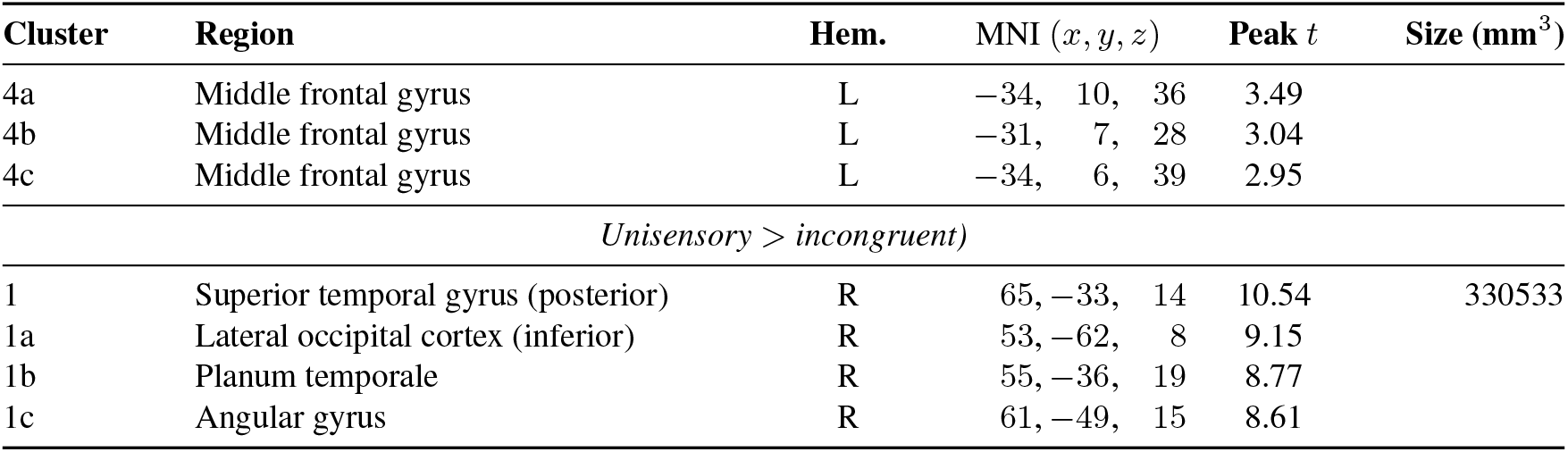
List of peak activations in the whole-brain RSA searchlight analysis. Significant clusters were identified using threshold-free cluster enhancement (TFCE) at *P <* 0.05. Bidirectional contrasts are reported in both directions (auditory vs. visual). Anatomical labels were assigned to each peak using the Harvard-Oxford atlas [71]. For each cluster, the peak and any local maxima at least 8 mm apart (labeled a, b, c) are shown. Clusters smaller than 20 voxels are omitted.

| Cluster | Region | Hem. | MNI ($x, y, z$) | Peak $t$ | Size (mm<sup>3</sup>) |
| --- | --- | --- | --- | --- | --- |
| *Auditory > visual* |  |  |  |  |  |
| 1 | Planum temporale | R | 59, -8, 2 | 13.22 | 30042 |
| 1a | Heschl's gyrus | R | 45, -24, 9 | 12.36 |  |
| 1b | Heschl's gyrus | R | 53, -14, 6 | 11.09 |  |
| 1c | Central opercular cortex | R | 50, -19, 12 | 10.36 |  |
| 2 | Heschl's gyrus | L | -51, -14, 2 | 12.96 | 19472 |
| 2a | Heschl's gyrus | L | -53, -21, 7 | 11.67 |  |
| 2b | Planum temporale | L | -58, -15, 4 | 10.69 |  |
| 2c | Heschl's gyrus | L | -45, -24, 7 | 10.46 |  |
| 3 | Superior frontal gyrus | R | 17, 23, 45 | 6.18 | 8041 |
| 3a | Superior frontal gyrus | L | -1, 37, 44 | 6.06 |  |
| 3b | Superior frontal gyrus | R | 9, 30, 44 | 5.69 |  |
| 3c | Superior frontal gyrus | L | -16, 27, 39 | 5.60 |  |
| 4 | Inferior frontal gyrus (pars opercularis) | R | 57, 18, 13 | 4.13 | 31 |
| 5 | Inferior frontal gyrus (pars triangularis) | R | 52, 25, 18 | 3.94 | 143 |
| *Visual > auditory* |  |  |  |  |  |
| 1 | Lingual gyrus | R | 26, -51, -8 | 19.21 | 530125 |
| 1a | Occipital fusiform gyrus | R | 32, -73, -8 | 18.82 |  |
| 1b | Occipital fusiform gyrus | R | 38, -67, -10 | 18.18 |  |
| 1c | Occipital fusiform gyrus | R | 28, -73, -7 | 17.80 |  |
| 2 | Temporal fusiform cortex (posterior) | R | 40, -10, -38 | 4.98 | 172 |
| 3 | Brain-stem | R | 16, -27, -33 | 4.15 | 180 |
| 4 | Unlabeled | L | -7, -62, -38 | 4.09 | 387 |
| 4a | Unlabeled | L | -14, -66, -37 | 2.86 |  |
| 4b | Unlabeled | L | -13, -58, -32 | 2.84 |  |
| 4c | Unlabeled | L | -8, -58, -43 | 2.76 |  |
| 5 | Temporal fusiform cortex (posterior) | R | 40, -9, -29 | 3.64 | 143 |
| 6 | Temporal fusiform cortex (anterior) | R | 34, -6, -36 | 3.40 | 60 |
| 7 | Brain-stem | R | 3, -16, -16 | 3.04 | 106 |
| 8 | Putamen | L | -22, 0, 7 | 3.03 | 40 |
| *Multisensory facilitation (congruent > auditory ∩ congruent > visual)* |  |  |  |  |  |
| 1 | Planum temporale | R | 62, -30, 19 | 9.12 | 24725 |
| 1a | Planum temporale | R | 51, -29, 14 | 8.13 |  |
| 1b | Planum temporale | R | 64, -21, 16 | 8.10 |  |
| 1c | Superior temporal gyrus (posterior) | R | 61, -31, 11 | 7.89 |  |
| 2 | Superior temporal gyrus (posterior) | L | -63, -36, 12 | 7.76 | 35829 |
| 2a | Parietal operculum cortex | L | -52, -33, 20 | 7.62 |  |
| 2b | Planum temporale | L | -60, -35, 12 | 7.33 |  |
| 2c | Parietal operculum cortex | L | -39, -35, 22 | 7.22 |  |
| 3 | Unlabeled | L | -33, -51, 9 | 3.67 | 255 |
| 4 | Insular cortex | R | 37, 14, -10 | 3.42 | 56 |
| 5 | Inferior frontal gyrus (pars opercularis) | R | 40, 16, 23 | 2.91 | 119 |
| 6 | Amygdala | L | -28, -3, -14 | 2.75 | 35 |
| 7 | Insular cortex | L | -39, 13, -9 | 2.68 | 32 |
| 8 | Temporal pole | R | 45, 12, -13 | 2.46 | 57 |
| *Crossmodal transfer* |  |  |  |  |  |
| 1 | Planum temporale | R | 59, -33, 14 | 9.92 | 237255 |
| 1a | Lateral occipital cortex (inferior) | R | 49, -61, 14 | 9.62 |  |
| 1b | Lateral occipital cortex (inferior) | R | 52, -65, 7 | 9.30 |  |
| 1c | Angular gyrus | R | 51, -58, 15 | 9.17 |  |
| 2 | Precentral gyrus | R | 48, 0, 47 | 6.56 | 324 |
| 3 | Superior frontal gyrus | L | -8, 1, 67 | 6.05 | 25 |
| 4 | Middle frontal gyrus | L | -32, 1, 40 | 3.66 | 629 |
| 4a | Middle frontal gyrus | L | -34, 10, 36 | 3.49 |  |
| 4b | Middle frontal gyrus | L | -31, 7, 28 | 3.04 |  |
| 4c | Middle frontal gyrus | L | -34, 6, 39 | 2.95 |  |
| *Unisensory > incongruent* |  |  |  |  |  |
| 1 | Superior temporal gyrus (posterior) | R | 65, -33, 14 | 10.54 | 330533 |
| 1a | Lateral occipital cortex (inferior) | R | 53, -62, 8 | 9.15 |  |
| 1b | Planum temporale | R | 55, -36, 19 | 8.77 |  |
| 1c | Angular gyrus | R | 61, -49, 15 | 8.61 |  |

